# *dlmoR:* An open-source R package for the dim-light melatonin onset (DLMO) hockey-stick method

**DOI:** 10.1101/2025.01.13.632603

**Authors:** Salma M. Thalji, Manuel Spitschan

**Affiliations:** Translational Sensory & Circadian Neuroscience, Max Planck Institute for Biological Cybernetics, Tübingen, Germany; Chronobiology & Health, TUM School of Medicine and Health, Technical University of Munich, Munich, Germany; TUM Institute for Advanced Study (TUM-IAS), Technical University of Munich, Garching, Germany

**Keywords:** Dim Light Melatonin Onset (DLMO), Circadian Rhythm, Biological Clocks, Sleep, Open Source Software, Algorithms

## Abstract

The dim-light melatonin onset (DLMO) is a commonly used circadian marker indicating the start time of evening melatonin synthesis in humans. Several quantitative techniques have been developed to determine DLMO from melatonin time series, including fixed- or variable-threshold techniques and the hockey-stick method developed by Danilenko et al. (2014). Here, we introduce *dlmoR*, an open-source (MIT License) implementation of the hockey-stick method written in R. Our clean-room implementation follows the original algorithm description, supported by iterative validation against the existing binary executable. We benchmarked *dlmoR* on 112 melatonin time series datasets from two independent studies (Heinrichs and Spitschan (2025); Blume et al. (2024)) and found high agreement with the reference implementation: mean discrepancies were −1.482 *±* 21.7 minutes for the Heinrichs and Spitschan (2025) dataset and 1.165 *±* 28.5 minutes for the Blume et al. (2024) dataset, with circular correlation coefficients of 0.964 and 0.986, respectively. Paired t-tests (*p* > 0.05) indicated no systematic difference or bias between methods. Beyond reproducing the hockey-stick algorithm, *dlmoR* adds capabilities absent from the original executable, including interactive visual diagnostics and bootstrapped confidence intervals, offering qualitative and quantitative views of estimation uncertainty. It supports programmatic, reproducible analysis of melatonin profiles, including batch processing and parameter manipulation. Leveraging this flexibility, we evaluated the sensitivity of the hockey-stick algorithm to controlled changes in sampling schedules, threshold levels, data completeness, and noise. Moderate changes, such as small timing jitter, limited data loss, or modest threshold shifts, kept estimates stable within *±*10 minutes, whereas pronounced alterations to sampling schedules, large multi-point deletions, or substantial threshold changes delayed estimates by over 40 minutes or prevented estimation. This analysis reveals fundamental limitations in the algorithm’s internal mechanics, particularly in how it identifies the onset window and models the melatonin rise, and underscores the need for new uncertainty-aware approaches to DLMO estimation.

## Introduction

The dim-light melatonin onset (DLMO) is the most reliable and widely used marker of circadian phase. By marking the onset of melatonin synthesis under dim light conditions in time, the DLMO provides precise insights into the synchronization of an individual’s circadian system with the 24-hour light/dark cycle (Pandi-Perumal et al. 2007; Keijzer et al. 2014). The DLMO is determined through bioanalytic assays of melatonin levels in saliva, plasma, or urine (Benloucif et al. 2008; Pandi-Perumal et al. 2007). The DLMO is instrumental in assessing phase delays, advances, and other deviations from expected circadian timing and plays a key role in research and clinical practice, aiding in the diagnosis and treatment of circadian rhythm sleep disorders, mood disorders and seasonal affective disorder (Pandi-Perumal et al. 2007; Keijzer et al. 2014).

Various methods have been developed to calculate DLMO, including fixed thresholds, dynamic thresholds, and visual estimation. However, these approaches have limitations: fixed thresholds may fail in low melatonin secretors, dynamic thresholds struggle with missing or inconsistent baseline data, and visual estimation is subjective and prone to variability (Kennaway 2023; Glacet et al. 2023). The hockey-stick algorithm (Danilenko et al. 2014) addresses these challenges by modeling melatonin rise as a piecewise linear-parabolic curve. This approach provides a more objective and reliable estimate of DLMO and has demonstrated superior accuracy, repeatability, and robustness compared to traditional methods (Danilenko et al. 2014; Glacet et al. 2023).

A Windows software implementation was developed and released by the authors (Danilenko and Verevkin 2020). However, its usability is limited by the constraints of being a binary executable. This closed-source format does not allow researchers to inspect, modify, or directly interact with the underlying implementation of the algorithm. Additionally, executable files without an API do not integrate seamlessly with widely used analytical workflows.

This paper introduces *dlmoR*, an open-source R package implementing the hockey-stick method. The package provides transparency into the algorithm, integrates seamlessly into analytical pipelines, and allows parameter adaptation for diverse experimental needs. Its open-source license fosters community contributions and ongoing improvements.

## Methods

### Algorithm analysis and prior implementation

Our open-source R implementation of the hockey-stick algorithm is based on our study of the original description by Danilenko et al. (2014), complemented by insights from personal communication with the author. A Windows executable of the algorithm, *hockeystickexe* (Danilenko and Verevkin 2020), is available as a free download but only in closed-source form. We adopted this executable as the reference against which our implementation was validated.

### Iterative development

Guided by the journal article and author correspondence, we created an initial R implementation (*dlmoR*) and iteratively compared its outputs with *hockeystickexe* on identical datasets. Each discrepancy highlighted aspects of the algorithm requiring adjustment. Through successive refinements, our implementation converged toward the behavior and objectives of the original algorithm.

### Implementation details

We chose to implement the hockey-stick algorithm in R (R Core Team 2024) due to its wide use, flexibility, open-source availability cross-platform support and modularity. Our implementation was designed in a modular fashion, with each module corresponding to a distinct component of the framework (see Fig. 1).

**Figure 1.**
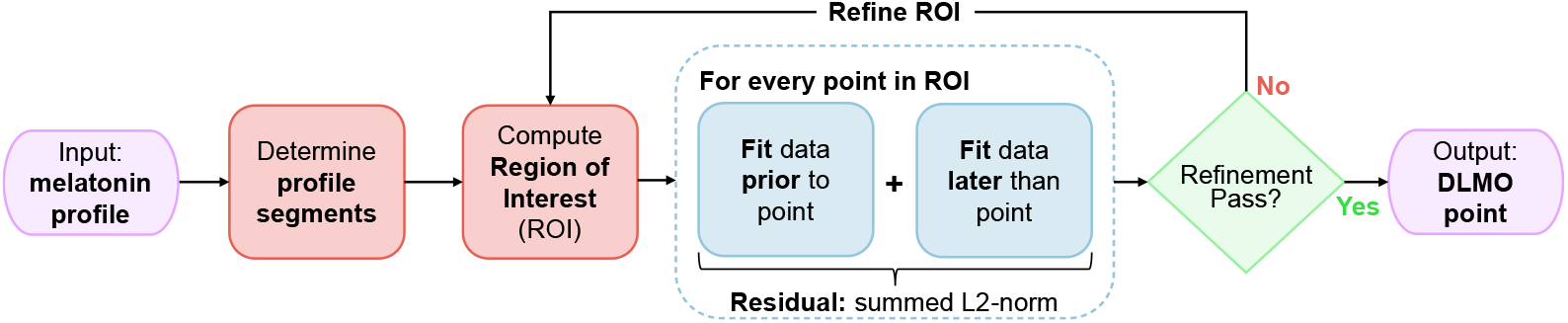
Flowchart of the *dlmoR* algorithm, which determines the dim light melatonin onset (DLMO) timestamp by identifying the inflection point in a melatonin profile. The algorithm first segments raw melatonin concentration data and defines a region of interest (ROI) for detecting melatonin rise. Candidate breakpoints within the ROI are evaluated in two phases: an initial coarse-grid pass across all points, followed by a fine-grid refinement phase restricted to the 10% with the lowest residuals from the initial pass. For each candidate, data before and after the point are fit separately, and residual error is computed as the summed L2-norm (Euclidean distance) between observed and fitted values. The point with the minimum residual from the refinement phase is identified as the DLMO point.xs

Each module was implemented as a separate script or set of scripts, focusing on the following key functionalities:

- *Preprocessing*: Functions handle the input data formatting and any required preprocessing steps, ensuring compatibility with the algorithm.
- *Core algorithm*: The core computation logic of the algorithm is encapsulated in dedicated modules, ensuring clarity and focus on the primary processing tasks.
- *Postprocessing and visualization*: Separate modules generate outputs, including tables and visualizations to aid in interpreting the results.

The implementation utilizes several R packages to streamline development and enhance functionality, including *dplyr* for data manipulation, *tidyverse* for efficient data handling, and *ggplot2* for visualization (Wickham et al. 2023, Wickham et al. 2019; Wickham 2016). Custom functions were developed where necessary to meet the specific requirements of the algorithm.

A comprehensive documentation system was embedded within the code, including inline comments and a user guide, to promote reproducibility and ease of use. Additionally, basic unit tests and validation checks were integrated into the workflow to ensure reliability and correctness. Error handling mechanisms, such as logging and exception handling, are also included to provide robust operational stability.

### Performance evaluation

To evaluate the performance of the *dlmoR* implementation, its concordance in dim-light melatonin onset (DLMO) estimation was assessed against the hockey-stick executable (*hockeystickexe*).

#### Datasets and sampling protocols

Two datasets comprising a total of 120 melatonin concentration time profiles were analyzed to determine DLMO using both methods.

- *Blume et al. (2024) dataset*: A total of 96 melatonin concentration time profiles, each consisting of 14 data points, were collected from 16 participants, each of whom attended three separate laboratory visits. Saliva samples (≥ 1 mL) were collected using Salivettes (Sarstedt, Nürnberg, Germany) at 30-minute intervals.
- *Heinrichs and Spitschan (2025) dataset*: A total of 24 melatonin concentration time profiles, each consisting of 23 data points, were collected from 12 participants during two-day laboratory visits. Each participant contributed two profiles, with saliva samples (≥ 1 mL) collected at 45-minute intervals.

The exact protocols and research questions asked in these two studies are irrelevant to the purpose of determining the DLMO, and are therefore not further described. In both datasets, melatonin concentrations were quantified using a radioimmunoassay (RIA, RK-DSM2, NovoLytiX GmbH, Switzerland) with a limit of quantification of 0.5–50 pg/mL, a detection limit of 0.2 pg/mL, a mean intra-assay precision of 7.9%, and a mean inter-assay precision of 9.8%.

#### Thresholds for DLMO analysis

For the performance evaluation, we assumed the following ascending level thresholds. For the Blume et al. (2024) dataset, an ascending level threshold of 5 pg/mL, consistent with the methodology outlined in Blume et al. (2024), was applied. For the Heinrichs and Spitschan (2025) dataset, the default ascending level threshold of 2.3 pg/mL was used.

#### Data exclusion

Eight of the 96 profiles in the Blume et al. (2024) dataset did not produce DLMO estimates using either method for one of three reasons: the profile was ambiguous, with no distinct baseline or discernible melatonin rise; the melatonin concentrations of the entire profile remained above the pre-defined threshold; or the entire profile remained below the pre-defined threshold. These data were excluded, leading to the inclusion of 88 time series from the Blume et al. (2024) dataset and 24 time series from the Heinrichs and Spitschan (2025) dataset, yielding a total of 112 datasets included in the analysis.

#### Analysis workflow

The analysis workflow differed between *dlmoR* and *hockeystickexe*. In *dlmoR*, melatonin profiles from both datasets were batch-processed using a custom script. The script parsed all melatonin profiles and determined the DLMO timestamp for each profile based on the user-defined ascending level threshold. Additionally, *dlmoR* saved a data structure for each profile containing the calculated DLMO timestamp, profile fit parameters, and the resulting plot. For analysis with *hockeystickexe*, each of the 112 melatonin profiles was manually entered into the *hockeystickexe* graphical user interface (GUI). The analysis for each profile was initiated by a button click and the resulting plot was saved manually. The DLMO timestamp printed in the GUI was manually tabulated in a spreadsheet alongside the corresponding sample name.

### Code, materials and data availability

All code, materials and data is available on our GitHub repository (https://github.com/tscnlab). *dlmoR* is available under the MIT License.

## Description of the hockey-stick algorithm

### Overall overview

The hockey-stick algorithm comprises three distinct phases (see Fig. 1). First, the input melatonin profile is validated and segmented into baseline and ascending intervals. Next, a region of interest (ROI) is established, within which the dim-light melatonin onset (DLMO) point will be sought in the subsequent phase. Finally, a piecewise function is fitted to the segmented melatonin profile, and the DLMO point is identified as the location within the ROI where the fitted profile yields the lowest residual. This time point is then reported as the output.

In the description that follows, we detail our interpretation of the algorithm, grounded in the original formulation by Danilenko et al. (2014), prior personal communication with the authors, and insights gained during the iterative development of our implementation. In this description, *node* refers to a melatonin data point, and *segment* refers to the region between a left node and a right node, including the right node but excluding the left node. More precise implementation details are provided as pseudocode in Appendix A.

The hockey-stick algorithm relies on several pre-defined parameter values—some documented in Danilenko et al. (2014), others not explicitly described. We reconstructed the rationale for these parameters through a close reading of the original description, informed by our correspondence with Prof. Danilenko and by systematic empirical testing during the development of *dlmoR*. This rationale is summarized in Table 1 of Appendix B. The open-source nature of *dlmoR* further allows advanced users to inspect and modify these parameters, enabling direct exploration of how such changes influence the DLMO estimation process.

**Table 1.** Overview of decision criteria used in the DLMO detection algorithm, organized by processing stage, with associated variables and physiological/algorithmic rationale.

| Algorithm Stage | Variable / Criterion | Rationale |
| --- | --- | --- |
| Ascending region definition —<br>Rules 2–3 | Steepness $\geq 1/2$ of the steepest segment (in non-base portion and preceding first ascending segment) | A relative steepness cut-off set at $1/2$ the maximum slope adapts to each profile's amplitude and rate of change. Applied near the start of the ascent, it avoids including slow, gradual rises that precede the main physiological melatonin increase. Because small, irregular slopes from measurement variability or short-term fluctuations are typically well below this relative threshold, they are also excluded as a side-effect. |
| Ascending region truncation —<br>Rule 1 | Zero or negative slope in final segment | Removes flat or falling tails from the ascending region, which cannot contribute to the rise and may bias the DLMO estimate, by altering the ascent model-fit, if retained. |
| Ascending region truncation —<br>Rule 2:<br>*Parallelogram rule* | Ratio of parallelogram's longest diagonal slope to lateral edge slope $> 1/2$ | The fitted parallelogram's lateral edge defines the steepest ascent that still encloses all ascending points and the last pre-ascending point (geometric upper bound), while the longest diagonal gives the average ascent over the region. Their ratio measures how closely the actual rise approaches the maximally steep, sustained rise allowed by the data. A ratio below $1/2$ indicates a brief, disproportionately fast late segment following a slow body, which can delay DLMO estimates; these are iteratively removed until the remaining ascent is both rapid on average and consistent across its span. |
| Ascending region truncation —<br>Rule 3 | Final segment $\geq 1/2$ the steepness of steepest remaining segment | After Rules 1 and 2 are applied, this rule trims late segments that rise too slowly. This prevents inclusion of gentle, tapering tails that contribute little to the rise but can shift the modeled DLMO later in time. |
| ROI definition | Line of interest: last 50% of final base segment (no intermediate segments) or from after last base node including all intermediate segments (if present) to first 95% of first ascending segment | Defines the time span used as the horizontal bounds of the ROI, targeting the transition from baseline to ascent. When no intermediate segments are present, starting at the last 50% of the final base segment captures the baseline portion closest to onset while avoiding distant flat baseline. When intermediate segments bridge the base and ascending phases, including all of them preserves transitional shapes, such as shallow early rises, needed for accurate onset detection. In both cases, ending at 95% of the first ascending segment avoids its terminal portion, where slopes may flatten, ensuring the ROI focuses on the part of the rise most representative of DLMO onset. |
| Coarse phase of DLMO search | Linear fit to earlier points constrained to pass through POI; slope limited to $-0.2$ to $0.2$ | Constraining the fit to pass through the POI ensures continuity between the two halves of the profile when estimating the inflection. Limiting the slope range to $[-0.2, 0.2]$ prevents unrealistic trends in the baseline portion: it avoids overly steep negative slopes that would imply implausible declines before melatonin onset, and overly steep positive slopes that could artificially inflate the apparent rate of rise, biasing the DLMO earlier. |
| Refinement phase of DLMO search | Refinement-phase parabola constraints (i–iv) | Passing through the POI (i) keeps the refined fit anchored to the candidate inflection point from the coarse phase. Ensuring the derivative is never zero (ii) avoids introducing local plateaus that would blur the definition of onset. Requiring the slope at the POI to be at least $1/2$ of the slope of the coarse-phase later linear fit (iii) prevents the refined curve from being unrealistically shallow at the onset compared to the initial coarse estimate. Demanding it be steeper than the coarse-phase earlier linear fit (iv) enforces a clear contrast between baseline and ascent. Together, these constraints refine the onset location while keeping the later fit both physiologically plausible and consistent with the data. |

### Validating the melatonin profile

First, the input melatonin profile is validated to ensure that it meets the necessary criteria for DLMO analysis using the hockey-stick method. Profiles are included in the analysis only if they exhibit a rise above a defined threshold (default value: 2.3 pg/mL, which is modifiable). If this criterion is not met, a warning is triggered, indicating the absence of a dynamic portion in the profile (“no dynamic part”). In addition, profiles with only one or two data points are considered insufficient for analysis and a corresponding warning is issued (“insufficient number of data points”). Finally, in cases where the profile shows a rise, but terminates with data points below the threshold, these terminal points are excluded from the analysis.

### Determining the profile segments

#### Defining the base region

Any segment below the threshold with a zero or negative slope (tan *θ* ≤ 0), or crossing the threshold in a downward direction, is classified as part of the *base* section. The *base* section also includes all segments located to the left of this segment. If a segment descends across the threshold or if the difference between the values of the two nearest nodes in the *base* section exceeds half the threshold value, the algorithm flags an inconsistency. In such cases, the following output is generated at the end of the module (“Profile consistency at base part: no”).

#### Defining the ascending region

The ascending region is defined as follows:

1. The segment that crosses the ascending threshold, along with all subsequent segments above the threshold, is classified as the *ascending* part. If the profile exhibits two rises above the threshold with an interval of less than two hours (default value), only the segments from the second rise onward are considered as part of the *ascending* section. The interval length is customizable and can be adjusted by the user.
2. In the portion of the profile not labeled as *base*, the *ascending* part also includes the steepest segment, all segments with a steepness exceeding half that of the steepest segment, and any segments situated between these steep segments. This rule does not exclude any segments identified in rule (1) but may add additional segments, including those below the threshold, if they meet the steepness criteria.
3. If the segment immediately preceding the first *ascending* segment (as determined by rules 1 and 2) has a steepness of at least half that of the *ascending* segment, it is also classified as part of the *ascending* section. Rule (3) is then reapplied iteratively to include additional segments if the same condition holds.

#### Truncating the ascending region

Three distinct rules are applied to truncate the *ascending* region, ensuring that the melatonin rise considered for determining the DLMO is both rapid and consistent.

1. First, the rightmost segment of this region is excluded from the *ascending* portion if it has a zero or negative slope (tan *θ* ≤ 0).
2. The parallelogram rule: After applying rule (1), segments of the *ascending* melatonin profile that rise too slowly are truncated using the properties of an optimal parallelogram fitted around all *ascending* points and the last data point preceding the *ascending* region (see Fig. 2). Specifically, the ratio of the slope of the parallelogram’s longest diagonal to the slope of its lateral edge must be greater than 1/2. If this condition is not met, the last segment of the *ascending* profile is truncated. This process is repeated iteratively, with the parallelogram being recalculated after each truncation, until the rule is satisfied. The construction of this parallelogram is subject to three constraints: (i) the lower edge must be horizontal and pass through the point with the lowest melatonin level, (ii) the upper edge must be horizontal and pass through the point with the highest melatonin level, and (iii) all *ascending* points must lie within the parallelogram. The term *optimal* refers to a parallelogram with the smallest possible area that satisfies these constraints. This construction and optimization process involves determining the lower-left x-coordinate of the parallelogram (*p*_*l*_), the lower-right x-coordinate (*p*_*r*_), and the slope of the lateral edge (*p*_*s*_) (see Fig. 2). By treating these as free parameters, the first and second constraints are inherently satisfied. The third constraint, that all *ascending* points are encompassed by the parallelogram, is treated as a soft constraint. Mathematically, the optimization can be expressed as follows:

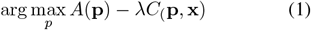

where **p** = (*p*_*l*_, *p*_*r*_, *p*_*s*_) is the optimization parameters, **x** are all datapoints labeled as *ascending* and the last point prior to the first *ascending* point, *A*(**p**) and *C*(**p, x**) are the functions that compute the area of the parallelogram and soft-constrained respectively, and *λ >>* 1 is a weighting scalar. We solve this optimization problem using the SANN algorithm available in the *Optim* package (Pan 2022) in R. For brevity, the formulation of *A*(**p**), *C*(**p, x**), and the initialization procedure are detailed in Appendix A, Algorithm 4.
3. The rightmost segment cannot be less than half as steep as the steepest of the remaining *ascending* segments (as defined after applying truncation Rules 1 and 2).

**Figure 2.**
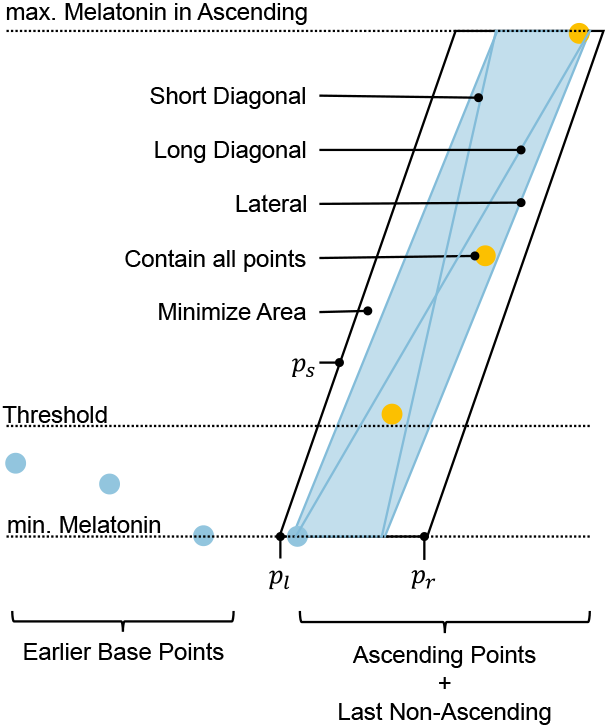
An illustration of the properties of a parallelogram used in the *parallelogram rule* for truncating the *ascending* region. The blue points represent *baseline* data, while the yellow points represent *ascending* data. The construction and optimization process involves determining the lower-left x-coordinate (*p*_*l*_), the lower-right x-coordinate (*p*_*r*_), and the slope of the lateral edge (*p*_*s*_). Annotations indicate the properties of an optimal parallelogram (i.e., the one with the smallest area) fitted around all ascending points and the last data point preceding the *ascending* segment.

Rules (2) and (3) can be disabled if desired.

#### Adding a segment to the left of the ascending region

If at this stage all segments belong to the *ascending* region, and the leftmost node is the only one that lies below half the threshold, an additional node is added 30 minutes to the left of it with a melatonin concentration level equal to either two times lower than the level of the leftmost node or zero, whichever is higher. The added segment is classified as *intermediate* (see below).

#### Truncating the base region

If the *base* part begins with nodes lying above the threshold, these nodes are excluded from the *base* part (see Fig. 4, Panel A). Although they remain visible in the figure, they are not included in the analysis.

#### Defining the intermediate region

Any segments lying between the *base* and *ascending* regions, that belong to neither of the two, are classified as *intermediate*.

### Computing the bounds of the ROI

The region of interest (ROI) refers to the portion of the melatonin profile that will be parsed in the subsequent stage of the algorithm to identify the DLMO point. First, a line of interest is defined for the left and right bounds of the ROI, and then the upper and lower bounds are defined.

#### Line of interest

To determine the line of interest, there are two scenarios:

1. *No intermediate point present*: If the *base* and *ascending* parts are defined, and no *intermediate* segments are present, the line of interest includes the last half (50%) of the final *base* segment and the first 95% of the first *ascending* segment. This interval is projected onto the time-axis and is denoted by a thick line parallel to the x-axis on the output plot (see Fig. 3). If there is only a single *base* node, the line of interest becomes narrower, spanning from this node to the first 95% of the first *ascending* segment.
2. *Intermediate points present*: If *intermediate* segments are present, the line of interest spans from immediately after the last *base* node, includes all *intermediate* segments, and extends to the first 95% of the first *ascending* segment.

**Figure 3.**
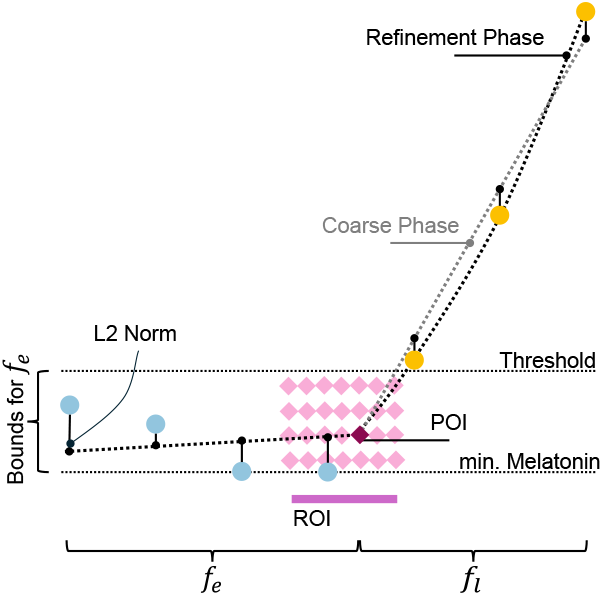
Schematic of the iterative grid search for the DLMO point. The *coarse phase* iterates over the region of interest (ROI), fitting piecewise functions to the *base* (*f*_*e*_) and *ascending* (*f*_*l*_) profile regions and calculating a composite residual for each point of interest (POI). The *refinement phase* narrows the ROI with finer resolution, and the POI with the lowest residual is identified as the DLMO time point.

**Figure 4.**
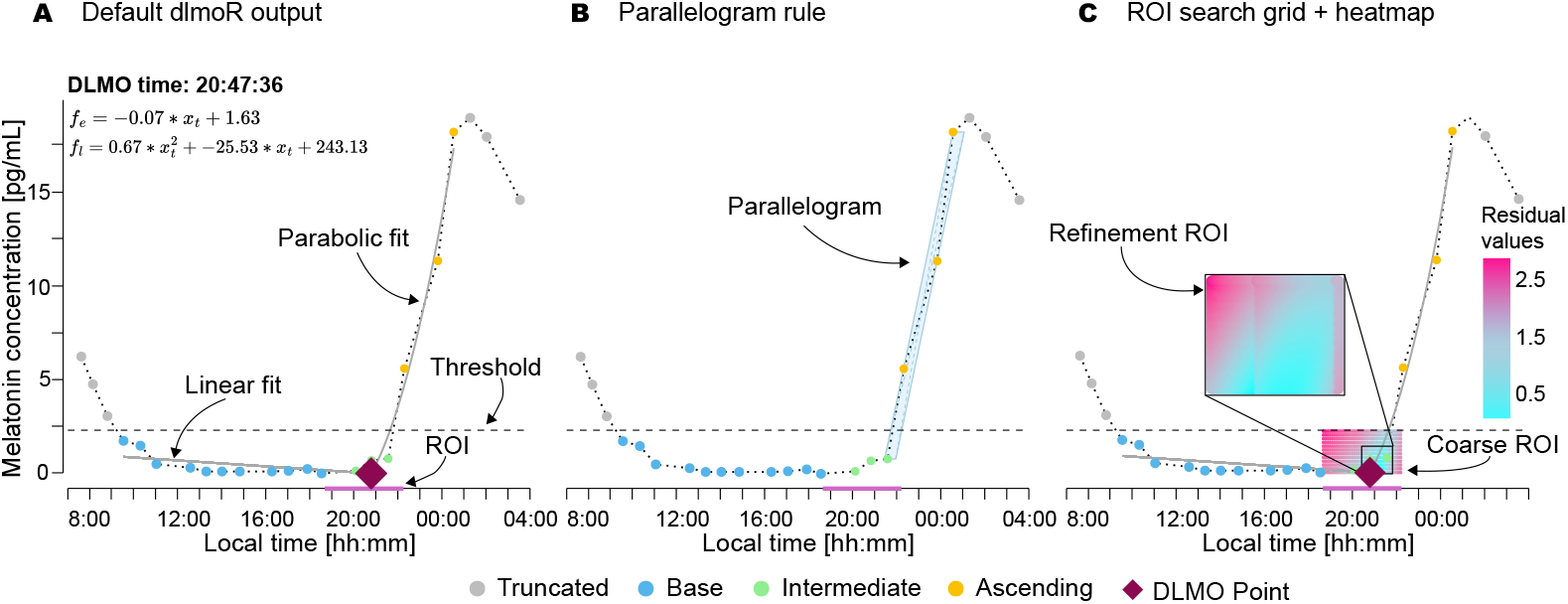
Visualizations of the hockey-stick method implemented in *dlmoR* with an example profile from the Heinrichs and Spitschan (2025) dataset. **A** Default *dlmoR* output, indicating various components of the fitted profile, including annotations of the linear (*f*_*e*_) and parabolic (*f*_*l*_) fit parameters. **B** Output including the visualization of the optimal parallelogram used for truncation of the ascending region of the profile. **C** Output with an overlay of the ROI fit residual heatmap used for DLMO point search.

#### ROI definition

The ROI is defined by the bounds of the line of interest, which set its left and right limits, while the upper and lower boundaries are determined by the threshold value and the melatonin level of the lowest node in the profile, respectively (see Fig. 3).

### Find the DLMO point

At this stage, all data points have been categorized into two (or three) groups: *base, ascending*, and (when present) *intermediate*. Additionally, an ROI has been defined within which we aim to identify the inflection point. Now, our method employs two phases to detect the inflection point: a coarse phase and a refinement phase (see Fig. 3).

#### Coarse phase of DLMO search

In the coarse phase, we perform a grid search over the ROI. The search uses step sizes of 0.1 in the time dimension (x-values) and 0.2 in the y-values. For each point of interest (POI) in the grid, the following steps are carried out:

1. *Linear fit to earlier points*: A linear model is fit that minimizes the *L*_2_-norm for all points occurring earlier than the POI, including the POI itself. The linear fit is constrained to pass through the POI, and its slope is limited to the range of −0.2 to 0.2. Additionally, the fitted line cannot exceed the highest y-value (threshold) or fall below the lowest y-value in the fitting domain.
2. *Linear fit to later points*: Similarly, a linear model is fit that minimizes the *L*_2_-norm for all points occurring later than the POI, including the POI itself. This line is also constrained to pass through the POI.

The overall cost associated with each POI is calculated as the sum of the residual *L*_2_-norms from the two linear fits.^*^

#### Refinement phase of DLMO search

In the refinement phase, we narrow the ROI to focus on the extrema of the 10% of points with the lowest costs from the coarse phase. This phase is similar in approach but uses finer step sizes of 0.01 in both dimensions to achieve greater precision (see Fig. 4).

During the refinement phase, a parabola is fit to the later points. This parabola must satisfy the following constraints: (i) the parabola is required to pass through the POI, (ii) the derivative of the parabola must never be zero, (iii) the slope of the parabola at the POI must be at least half as steep as the slope of the linear fit identified during the coarse phase and (iv) steeper than the slope of the linear fit for the earlier points.

Generally, for both the coarse and refinement phases, the cost of a point can be expressed as:

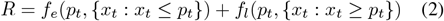

where *x*_*t*_ represents all data points and *p*_*t*_ is the point of interest (POI). Specifically:

- {*x*_*t*_ : *x*_*t*_ ≤ *p*_*t*_} denotes all points earlier than and/or equal to the POI.
- {*x*_*t*_ : *x*_*t*_ ≥ *p*_*t*_} denotes all points later than and/or equal to the POI.

Here, *f*_*e*_ is the linear fit function applied to the earlier data points, and *f*_*l*_ is the fit function for the later data points, which can be either linear or parabolic depending on the search phase. Details of this implementation can be found in Appendix A, Algorithm 9.

This two-phase design, coarse search followed by refinement, follows the original hockey-stick algorithm described by Danilenko et al. (2014), which uses the coarse phase to efficiently locate a promising ROI before applying a finer grid to that reduced region. While the refinement step is more computationally demanding, it is still substantially faster than applying a fine grid across the entire parameter space. In *dlmoR*, both coarse and refined results are retained and reported, whereas *hockeystickexe* outputs only the final refined estimate. This modularity allows *dlmoR* users to run only the coarse phase if a faster, approximate estimate is sufficient (typically under one minute per profile), or to include the refinement phase for maximum precision (typically 2–3 minutes per profile), with exact runtimes depending on ROI size and hardware performance.

## Results

### Independent implementation of the hockey-stick algorithm

Despite the original hockey-stick software being only available as a closed-source Windows executable, we were able to derive an R-based implementation of the hockey-stick algorithm developed by (Danilenko et al. 2014).

### Performance evaluation

We assessed the concordance in dim light melatonin onset (DLMO) time estimates derived from the two methods – *dlmoR* and *hockeystickexe* – applied to two datasets comprising a total of 120 melatonin time series Blume et al. (2024); Heinrichs and Spitschan (2025), with a total of 112 usable datasets. Circular statistics were employed to evaluate concordance due to the periodic nature of time data.

#### Descriptive Statistics

The circular difference in DLMO time estimates between the two methods were summarized using descriptive statistics, namely the mean and standard deviation, to describe the central tendency and dispersion. The mean difference in DLMO estimates between *dlmoR* and *hockeystickexe* is 1.165 *±* 28.5 minutes for the Blume et al. (2024) dataset, and −1.482 *±* 21.7 minutes for the Heinrichs and Spitschan (2025) dataset.

#### Outliers

Outliers were identified by manually inspecting data profiles with DLMO estimate discrepancies between methods exceeding *±*1 hour. Of the 112 profiles analyzed, two significant outliers were observed: a DLMO estimate difference of approximately +80 minutes and one of approximately +240 minutes, indicating that the *dlmoR* method estimated DLMO timestamps more than one hour later than *hockeystickexe*. Close inspection of each scenario highlighted two key scenarios, illustrated in Appendix C in which *dlmoR* and *hockeystickexe* analysis differs: (1) the handling of spurious above-threshold melatonin rises occuring early in the time profiles, and (2) differing interpretrations of multiple rises above threshold between methods. In the case of the melatonin profile with a between-method DLMO difference of 80 minutes, a single melatonin sample briefly rose above the threshold, followed by a decline and a subsequent steady rise within one hour (Case 1, see Appendix C, Fig. 13, Panel A). The *hockeystickexe* algorithm fit the earlier single-point rise, whereas *dlmoR* fit the later steady consecutive rise. For the melatonin profile with a +240 minute difference, a spurious single-point rise near the beginning of the profile (second point in a 14-point time series) was fit as the DLMO by *hockeystickexe. dlmoR* disregarded this as noise and fit the steady rise occurring four hours later (Case 2, see Appendix C, Fig. 13, Panel B). Without outliers, the mean discrepancy in DLMO estimate between methods was −2.082 *±* 11.1 minutes for the Blume et al. (2024) dataset. The small mean differences in both datasets suggest minimal overall difference in DLMO estimate between the methods. The larger standard deviation in the Blume et al. (2024) dataset reflects greater variability in the differences compared to the Heinrichs and Spitschan (2025) dataset.

#### Method agreement within specific time thresholds

The agreement between methods was further assessed by categorizing DLMO estimates based on time difference thresholds. For the Blume dataset, 46 estimates (52.3 %) were within *±*5 minutes of each other, 69 estimates (78.4%) were within *±*15 minutes, and the difference in three estimates (3.41%) exceeded *±*30 minutes. For the Heinrichs dataset, 16 estimates (66.7%) were within *±*5 minutes, *±*18 estimates (75%) were within *±*15 minutes, and the difference in 4 estimates (16.7%) exceeded *±*30 minutes (see Fig. 5, Panel B).

**Figure 5.**
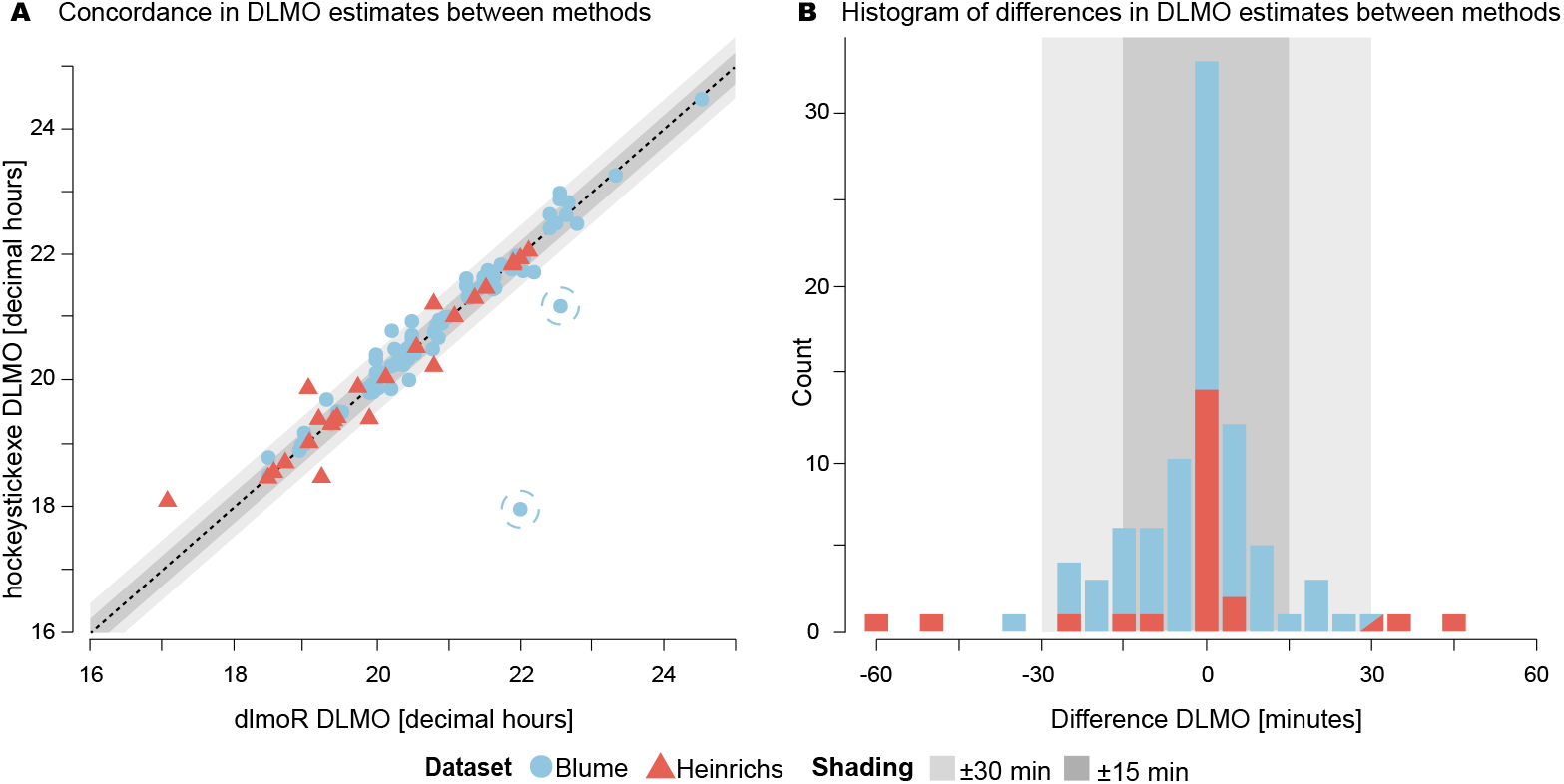
Performance evaluation of *dlmoR* against *hockeystickexe*. **A** Concordance in DLMO estimate between implementations, with two outliers highlighted by circles (see subsection *Outliers* for details and Appendix C for visualization). **B** Histogram of differences in DLMO estimates between methods, with outliers removed.

#### Concordance and bias detection

Circular correlation coefficients and paired t-tests were used to evaluate the strength and significance of agreement and to identify systematic biases between the two methods. Circular correlation values were 0.986 (0.911 with outliers) for the Blume et al. (2024) dataset and 0.964 for the Heinrichs and Spitschan (2025) dataset, indicating strong agreement between the methods in both datasets (see Fig. 5, Panel A). Circular paired t-tests yielded p-values of 0.088 (0.601 with outliers) for the Blume dataset and 0.742 for the Heinrichs dataset, suggesting no systematic difference or bias between the methods.

#### Skewness and bias

Analysis of skewness revealed minor biases in DLMO estimation. The Blume et al. (2024) dataset had a skewness of −0.22 (6.37 with outliers), while the Heinrichs and Spitschan (2025) dataset had a skewness of −0.796. These values suggest that *dlmoR* tends to provide slightly earlier estimates compared to *hockeystickexe*.

#### Visual inspection and residual analysis

Visual inspection of fitted melatonin profiles revealed two primary contributors of discrepancies between methods, aside from identified outliers:

1. *Parallelogram-based ascending segment truncation*: The methods differed in the truncation of *ascending* segments, leading to the fitting of different sets of points. This impacted the distribution of residual heatmaps and the selection of the DLMO point, which corresponds to the timepoint with the lowest fit residual.
2. *Diffuse region of interest (ROI) heatmap*: The *dlmoR* visualization indicated diffuse ROI heatmaps with multiple points offering similarly good fits (differences on the order of 1 *×* 10^−2^). Minor variations in fit procedures likely contributed to residual variation. When small deviations in the fits lead to minor differences in the residuals, a diffuse residual heatmap can correspond to differences in DLMO estimates of 15 minutes or more (see example in Fig. 4, Panel C and subsection *The residual heatmap* below for details).

### R package

*dlmoR* is an R package developed to provide a streamlined and efficient tool for analyzing dim light melatonin onset. The package is designed to handle both individual and batch processing of data profiles and offers a range of customizable features to meet diverse user needs.

#### Key functionalities of the dlmoR package

The core functionalities of the *dlmoR* package are:

- *Data visualization*: Generates detailed plots that include the following features (see Fig. 4):
  - The raw data profile
  - The subset of points analyzed
  - The identified DLMO timestamp (labeled on the plot and output as an annotation)
  - The two segments of the hockey-stick fit with annotations of their corresponding parameters
  - A time segment highlighting the region of interest for inflection point detection
  - A residual heatmap for points within the region of interest grid, providing insight into the confidence or definitiveness of the result
  - A parallelogram indicating the decision boundary used to trim the points of melatonin rise, offering a clear understanding of how the subset of rise points for the fit was determined
- *Customizability*: Allows users to:
  - Set the ascending threshold
  - Set the multi-rise interval length
  - Modify plot features for tailored visualization
- *Batch processing*: Supports the analysis of multiple data profiles in a single run, enhancing efficiency for large datasets.
- *Automated Outputs*: Automatically saves all key results, including:
  - The raw data profile
  - Labeled segments
  - Fit parameters
  - DLMO point
  - Residual heatmap
  - The generated visualization plot

#### The residual heatmap

The residual heatmap spans the full region of interest (ROI) explored during the incremental search for the DLMO—the point of lowest model-fit residual. Both its bounds and values are algorithmically determined. The range is bounded below by the lowest melatonin value in the profile and above by the threshold value, while the domain bounds are defined relative to the labeled regions of the melatonin profile (*baseline, intermediate, ascending*). These regional labels are applied according to the rules described by Danilenko et al. (2014) in the original hockey-stick algorithm and depend on both the threshold value and the inter-point slopes of melatonin concentrations in the time profile. The values in the heatmap represent the residual of the hockey-stick model fit at each point in the grid. Because the heatmap directly represents this residual search space, it cannot be arbitrarily modified by the user. Instead, it serves as a diagnostic visualization of how the algorithm evaluates candidate DLMO points within the ROI.

While *hockeystickexe* produces a single DLMO estimate that gives the impression of an absolute, definitive solution, *dlmoR* exposes this underlying residual landscape and thereby uncovers a plurality of plausible solutions. Sharp, isolated minima indicate a well-constrained estimate, whereas diffuse or multi-modal minima reveal that the solution is embedded within a landscape of alternatives rather than being singular or definitive. In practice, many candidate estimates can yield very similar residuals while differing substantially in clock time. For instance, in the example showcased in Fig. 6, the top 0.4% of residuals (31 out of 10,220 grid points) span only 0.008 in residual value (0.123–0.131) yet correspond to DLMO estimates 31.8 minutes apart.

**Figure 6.**
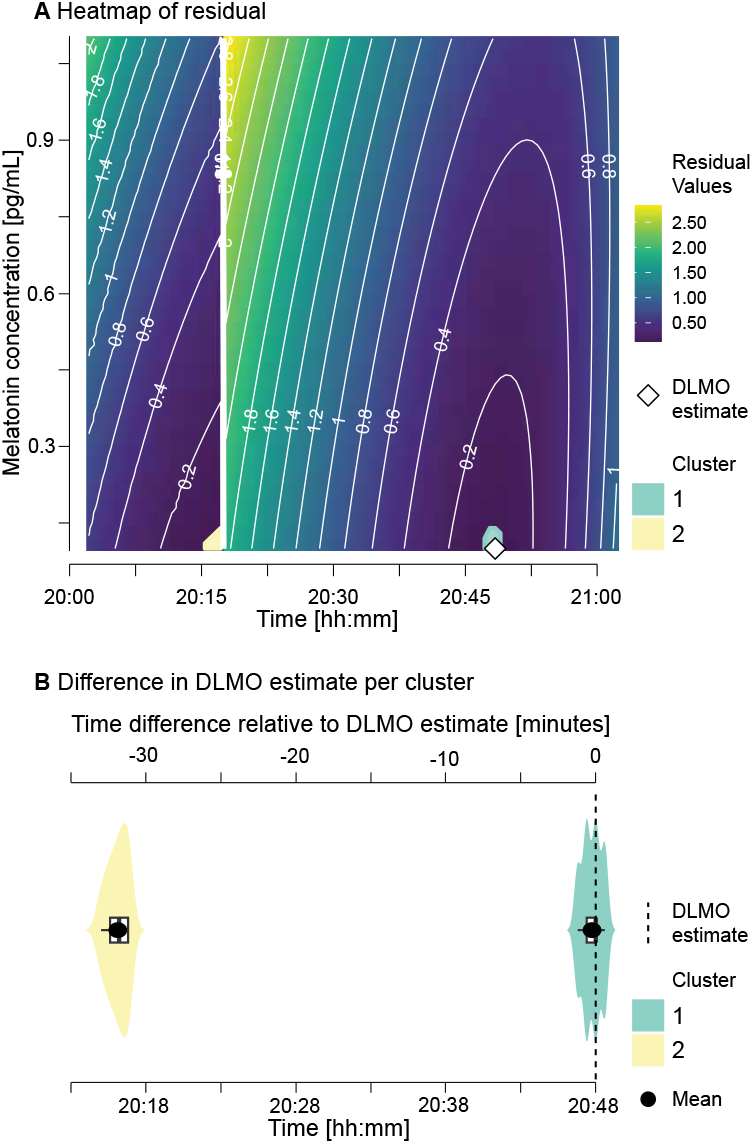
Residual landscape of candidate DLMO estimates. **A** Heatmap of hockey-stick model fit residuals for all grid points in the region of interest (ROI) for an example melatonin profile, with colors indicating residual value and contour lines tracing residual magnitude across the landscape. In this example, colored clusters mark grid points within the 0.4% lowest residuals. The *dlmoR* estimate (white diamond), corresponding to the point with the lowest residual, lies within cluster 1, while cluster 2, occurring earlier in time, contains residual values of similar magnitude, illustrating that multiple low-residual regions can exist. **B** Clock times of candidate estimates in each cluster as violin plots relative to the DLMO estimate.

To make these landscapes accessible, we developed a Shiny app for interactive diagnostics in *dlmoR* (Appendix D, Fig. 14). The *Residual Heatmap Explorer* extends static residual heatmaps into an interactive tool, allowing examination of clusters of good model fit and assessment of alternative minima through distributional and statistical summaries. This interactivity reveals the hidden multiplicity of solutions, clarifying how sharply or diffusely candidate estimates are defined and generalizing the illustrative example in Fig. 6 into a reusable diagnostic.

#### DLMO estimate confidence intervals

In addition to the qualitative lens offered by the heatmap exploration, *dlmoR* provides quantitative tools to formalize uncertainty assessment. We have implemented four complementary bootstrap methods for inference of DLMO confidence intervals: (i) *Monte Carlo bootstrap*, which adds Gaussian jitter to sampling times and melatonin concentrations to simulate experimental such as protocol delays and assay noise; (ii) *residual bootstrap*, which resamples model residuals to test robustness of the fitted hockey-stick model; (iii) *wild bootstrap*, which preserves heteroscedasticity through random multipliers, making it suitable for datasets with uneven measurement precision; and (iv) *hybrid bootstrap*, which combines time jittering with residual resampling, capturing both external noise and model-fit variability in a single procedure. Each method generates an empirical distribution of DLMO estimates, summarized by its mean and an empirical 95% confidence interval. In addition, a histogram of bootstrapped DLMO estimates is generated, annotated with the original estimate, the bootstrap mean and the 95% confidence interval bounds.

Together with the residual heatmaps and the Shiny app, these bootstraps ensure that *dlmoR* does not obscure uncertainty behind a single estimate but instead makes visible and quantifiable the landscape of plausible alternatives. A comparison of the different bootstrap methods, the noise sources they simulate, and the typical use cases for each is provided in Appendix E.1, Table 2. Example outputs from each bootstrap method are shown in Figure 15 of Appendix E.2.

**Table 2.** Comparison of bootstrap methods implemented in *dlmoR*.

| Method | Noise source(s) simulated | Key parameters | Typical use case |
| --- | --- | --- | --- |
| **Monte Carlo** | Simulates realistic experimental variability by perturbing both sampling times and melatonin concentrations:<br>Temporal jitter of samples<br>Heteroscedastic assay noise proportional to concentration | *time_sd*: SD of time jitter (default = $\frac{1}{2}$ minimum sampling interval)<br>*mel_cv*: coefficient of variation for assay noise (default = 0.079)<br>*clip_negatives*: enforce non-negative concentrations | When protocol timing errors (late/early sampling) and assay variability substantially contribute to DLMO uncertainty; realistic simulation of experimental conditions |
| **Residual** | Quantifies variability due to deviations around the fitted hockey-stick model without introducing timing or assay noise:<br>Resampling residuals from baseline and ascending fits | *clip_negatives*: enforce non-negative concentrations | When timing and assay precision are well-controlled; isolates uncertainty from measurement scatter around the model fit |
| **Wild** | Generalizes the residual bootstrap by applying random weights to residuals, preserving heteroscedasticity:<br>*rademacher*: $\pm 1$ sign flips<br>*normal*: Gaussian weights | *wild_type*: residual weighting scheme<br>*clip_negatives*: enforce non-negative concentrations | Robust uncertainty estimation under heteroscedastic residuals; *rademacher* ($w \in \{-1, +1\}$) for robustness, *normal* ($w \sim \mathcal{N}(0, 1)$) for Gaussian-like variability |
| **Hybrid** | Combines Monte Carlo and residual approaches to simulate both timing irregularities and residual noise:<br>Temporal jitter of samples<br>Residual resampling added to jittered model | *time_sd*: SD of time jitter (default = $\frac{1}{2}$ minimum sampling interval)<br>*clip_negatives*: enforce non-negative concentrations | Conservative choice for sensitivity analysis; captures both experimental (timing) and statistical (residual) uncertainties |

To illustrate how *dlmoR* extends and enhances the original *hockeystickexe* implementation, we provide a side-by-side comparison in Appendix F, Table 3. The table outlines key differences in platform, reproducibility, diagnostics, uncertainty quantification, and extensibility. In particular, it highlights the advances provided by *dlmoR*, including (i) open-source, cross-platform implementation in R; (ii) full parameter customization (e.g., threshold levels, base slope limits, multi-rise interval lengths); (iii) customizable plotting, batch processing, and scripting support; (iv) reproducible workflows with transparent implementation of the hockey-stick algorithm; and two analytical functionalities not available in the original executable: (v) visualization of model-fit residual heatmaps with an accompanying Shiny app for interactive exploration; and (vi) bootstrap-based methods for formal uncertainty quantification of DLMO estimates.

**Table 3.** Comparison of dlmoR and hockeystickexe feature availability.

| Feature | *hockeystickexe* | *dlmoR* |
| --- | --- | --- |
| **Platform** | • Windows-only executable<br>• Requires virtualization or compatibility layer to run on macOS/Linux | • Runs natively on Windows, macOS, and Linux via R<br>• Works in local, server, and cloud environments |
| **License** | Closed source (unknown) | Open source (MIT License) |
| **Residual Heatmap** | Not provided | • Provided numerically as residual matrices (coarse and fine)<br>• Visual overlay in plots<br>• Interactive Shiny app for zooming and stats |
| **Estimate Confidence Intervals** | Not provided | Five bootstrapping methods for *post hoc* CI estimation |
| **Batch Processing** | Not supported | Fully supported via scripting |
| **Parameter Customization** | • Threshold<br>• Base segment slope limits | • Threshold<br>• Base segment slope limits<br>• Multi-rise interval length<br>• Optional coarse-only fitting mode<br>• Full access to algorithm internals (open source) |
| **Plotting** | • Fixed-format PNG<br>• No customization | • Export formats: PNG, JPEG, SVG, R object<br>• Fully customizable:<br>• Melatonin profile<br>• Model fit overlays<br>• DLMO estimate<br>• ROI and threshold lines<br>• Segment classification<br>• Residual heatmaps (coarse and fine)<br>• Trimming parallelogram overlay |
| **Outputs** | • DLMO in HH:MM:SS<br>• Annotated plot of fine phase fit | • DLMO in HH:MM:SS, POSIXct, decimal hours<br>• Coarse and fine model parameters<br>• Grid intercepts and residuals<br>• Full plot object for coarse and fine phases<br>• Complete data structure (fully scriptable) |
| **Output Saving Options** | • PNG only<br>• Fixed file path and name | • Fully scriptable saving<br>• Directory control and custom naming<br>• Multiple formats<br>• Selective component output |

### In-depth investigation of hockey-stick algorithm behavior

Having demonstrated that *dlmoR* sufficiently concords with the *hockeystick*.*exe* implementation, we next leveraged its flexibility to examine the robustness and sensitivity of DLMO estimates under diverse conditions. By programmatically modifying both analysis parameters and melatonin time series characteristics, we conducted five sets of analyses designed to probe how robustly the hockey-stick algorithm recovers the DLMO timing across different scenarios. In particular we investigated the effects of:

- **Varying sampling frequency**: to assess how changes in temporal resolution and the number of available data points influence the DLMO estimate.
- **Altering threshold values**: to determine how different choices of melatonin threshold affect the resulting DLMO estimate.
- **Irregular sampling and noise perturbations**: comprising (i) single-point deletions, (ii) multi-point deletions, and (iii) added noise to melatonin concentration or sample timing, to test robustness under realistic data imperfections.

We expand on the rationale, methods, and results of each analysis in the sections below.

#### Computational environment

All computational analyses detailed below were performed either on a MacBook Pro with Apple M4 Max (12-core CPU, 38-core GPU, 16-core Neural Engine) and 36 GB unified memory, or on the NYX high-performance computing (HPC) cluster at the Max Planck Institute for Biological Cybernetics. Each NYX compute node is equipped with two AMD EPYC 7452 32-core processors (2.35 GHz; 128 logical CPUs total), and jobs were run with exclusive node access. Wall-clock times and corresponding core-hour usage for the analyses are reported in Appendix G.

#### Sampling frequency

##### Rationale

This analysis investigates the sensitivity of DLMO estimates to temporal resolution of the melatonin time series, considering both dense and sparse sampling schemes. Reduced sampling rates are particularly relevant in applied research settings, where melatonin assays are costly and frequent sampling entails substantial logistical effort. Understanding how DLMO estimates behave under sparse sampling can inform practical study design decisions.

##### Methods

To quantify this effect, we performed a resampling analysis on the Blume et al. (2024) and Heinrichs and Spitschan (2025) datasets. For each melatonin profile, we used *dlmoR* to generate a reference DLMO estimate. We then resampled the profile using piecewise cubic Hermite interpolation (PCHIP), which preserves the shape and monotonicity of the original signal while avoiding spurious oscillations. Resampled time series were generated at intervals of 2, 5, 10, 15, 20, 30, 45, 60, 75, and 90 minutes, simulating progressively coarser sampling schemes. DLMO was re-estimated at each interval, and differences from the reference DLMO were computed to assess sensitivity to temporal resolution.

For the Blume et al. (2024) dataset, sampling times were assigned based on the nominal 30-minute protocol schedule, and the original DLMO estimates were calculated using these idealized timestamps. In contrast, the Heinrichs and Spitschan (2025) dataset included actual recorded sample times, which varied slightly from the nominal 45-minute protocol due to real-world procedural deviations. As a result, the reference DLMO estimate in Heinrichs and Spitschan (2025) did not align exactly with any fixed resampling interval.

##### Results

Across both Heinrichs and Spitschan (2025) and Blume et al. (2024) datasets, DLMO estimates exhibited varying sensitivity to sampling interval, with differences in both magnitude and pattern (see Fig. 7 and Fig. 8 as well as Appendix H.1 for Tbl. 6 and Tbl. 7).

**Figure 7.**
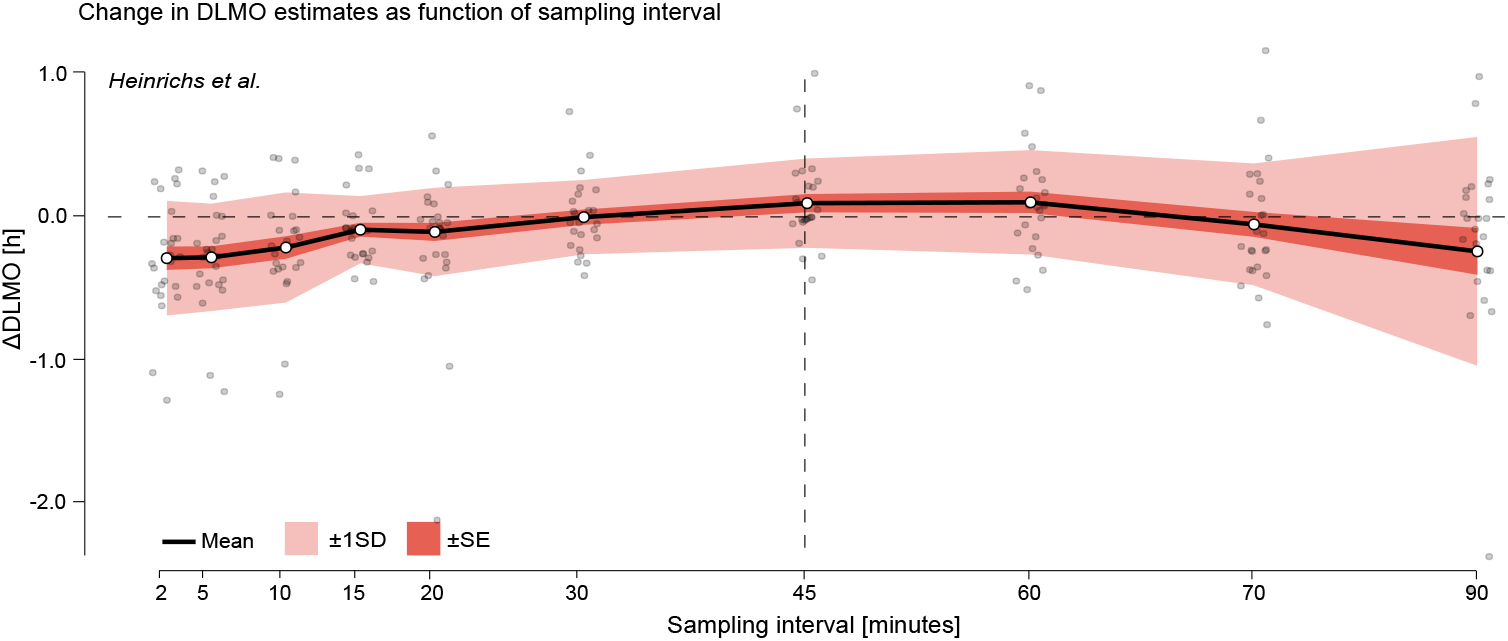
Change in DLMO estimates (in decimal hours) as a function of sampling interval for the Heinrichs and Spitschan (2025) dataset. Black line shows the mean across all melatonin profiles of the difference between the DLMO estimated at each sampling interval and the reference DLMO, calculated from the actual recorded sample times, which deviated from the nominal 45-minute protocol. Shaded regions indicate variability across participants: dark shading shows *±*1 standard error (SE), and lighter shading shows *±*1 standard deviation (SD). Grey points represent individual melatonin profiles at each resampled interval. The vertical dashed line marks the nominal 45-minute sampling interval of the study protocol.

**Figure 8.**
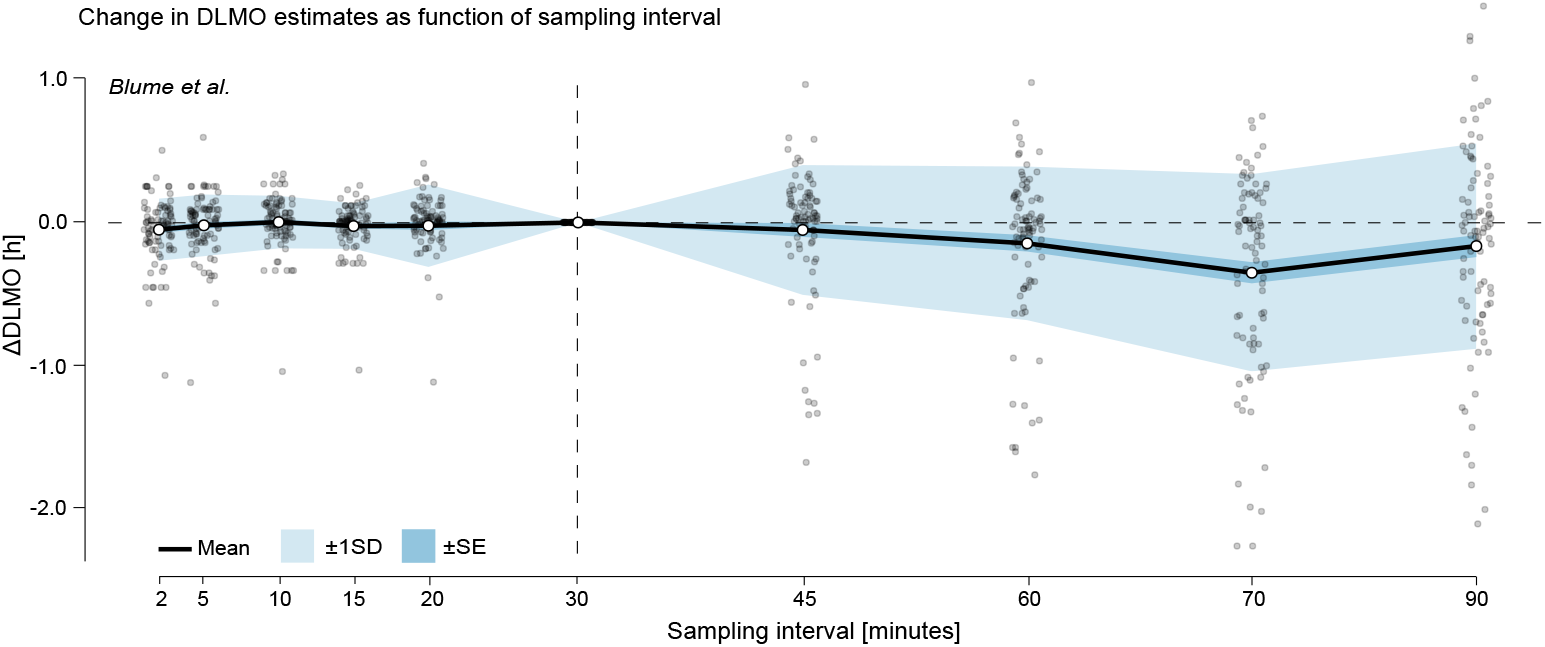
Change in DLMO estimates (in decimal hours) as a function of sampling interval for the Blume et al. (2024) dataset. Black line shows the mean across all melatonin profiles of the difference between the DLMO estimated at each sampling interval and the reference DLMO, calculated from the actual recorded sample times, which followed the nominal 30-minute protocol. Shaded regions indicate variability across participants: dark shading shows *±*1 standard error (SE), and lighter shading shows *±*1 standard deviation (SD). Grey points represent individual melatonin profiles at each resampled interval. The vertical dashed line marks the nominal 30-minute sampling interval of the original protocol.

In the Heinrichs and Spitschan (2025) dataset, both the mean deviation and standard deviation (SD) were elevated at short intervals (e.g., at 2–minutes: mean ΔDLMO =− 0.2522 *±* 0.3540 hours), then decreased at moderate sampling intervals (15–60 minutes), reaching a minimum SD of 0.2295 hours at 30–minute sampling (see Fig. 7 and Appendix H.1, Tbl. 6). At coarser intervals (75– and 90– minute), both mean deviation and SD increased again, with SD rising to 0.3765 hours and 0.7073 hours, respectively. This non-monotonic pattern, with elevated variability at both fine and coarse intervals and lowest variability at intermediate resolutions (e.g., 30–45 minutes), suggests that the relationship between sampling interval and DLMO stability is not strictly linear in this dataset.

Despite the study’s nominal 45–minute sampling interval, the deviation at 45–minutes sampling interval was not zero (mean ΔDLMO = 0.0877 hours). This reflects that the original DLMO estimates were based on the actual sample timestamps, which varied slightly due to protocol time deviations.

In the Blume et al. (2024) dataset, the mean deviation remained close to zero for intervals up to 30–minutes (e.g., mean ΔDLMO at 20–minutes= ≤0.0226 hours), and SD stayed ≤0.2853 hours (see Fig. 8 and Appendix H.1, Tbl. 7). At 30–minutes, no deviation was observed across all profiles because the dataset used nominal timestamps based on the protocol’s fixed 30–minute sampling schedule, which aligned exactly with the 30–minute resampling interval. At longer intervals (60–90–minutes), variability and downward bias increased, with the largest mean shift at 75–minutes (−0.3486 hours) and SD peaking at 90–minute sampling intervals (−0.7119 hours). Note that for this dataset, some resampled profiles did not yield a DLMO estimate at certain intervals, as they remained entirely sub-threshold or supra-threshold and therefore lacked the threshold crossing required by the hockey-stick algorithm.

Overall, DLMO estimates were relatively stable for intervals of ≤30–minutes, while intervals of ≥60–minutes introduced substantial variability and, in some cases, systematic bias in the mean DLMO time.

#### Threshold values

##### Rationale

Another key factor in DLMO estimation is the choice of melatonin threshold. The hockey-stick algorithm requires defining a concentration that marks the transition from baseline to rise, yet no consensus exists on which value to use. Because thresholds are often chosen arbitrarily or vary between studies, it is important to quantify how sensitive DLMO estimates are to this parameter.

##### Methods

DLMO was estimated for each profile at thresholds of 2, 3, 4, 5, and 10 pg/mL, and shifts in DLMO timing were calculated relative to the 2 pg/mL estimate.

##### Results

Increasing the threshold systematically delayed the estimated DLMO, with progressively larger shifts, relative to the 2 pg/mL estimate, at higher thresholds (see Fig. 9 and Appendix H.2 for Tbl. 10 and Tbl. 11). In the Blume et al. (2024) dataset, mean shifts increased from 0.2847 hours at 3 pg/mL to 0.97110 hours at 10 pg/mL, with variability (SD) rising from 0.5856 hour to 1.1041 hour. In contrast, the Heinrichs and Spitschan (2025) dataset showed smaller overall shifts (e.g., mean = 0.1067 hours*±*0.2048 hours at 10 pg/mL). Lower thresholds (2–4 pg/mL) produced estimates that were more closely aligned, with SDs remaining below 0.12 hours.

**Figure 9.**
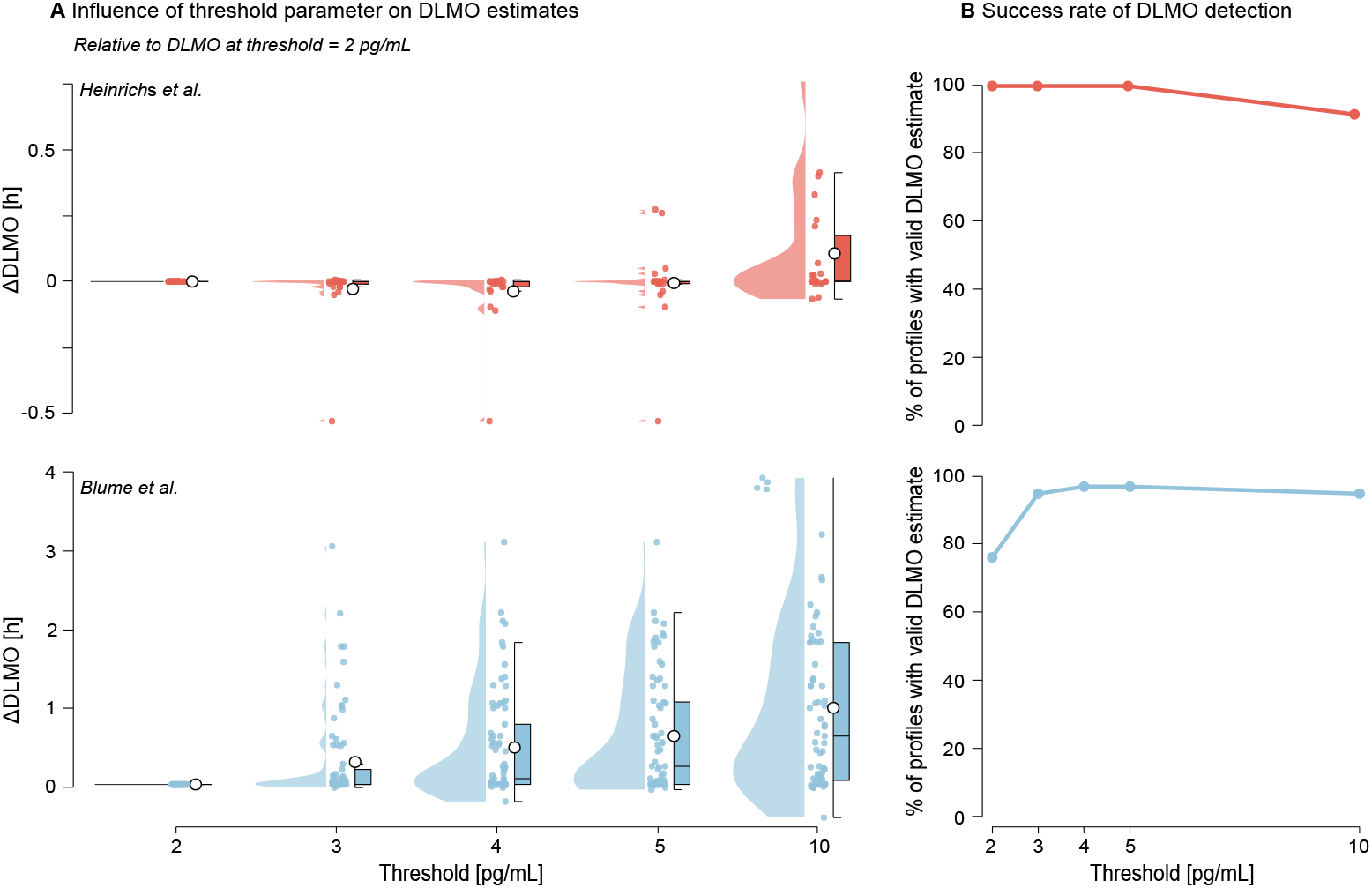
Effect of melatonin concentration threshold value on DLMO estimates and detection success for the Heinrichs and Spitschan (2025) (red) and Blume et al. (2024) (blue) datasets. **A** Change in DLMO estimates (in decimal hours) at each threshold, relative to the estimate at a 2 pg/mL threshold. Points represent individual profiles, with boxplots indicating the median and interquartile range and violins showing the distribution. Higher thresholds generally delayed the estimated DLMO, with larger shifts in the Blume et al. (2024) dataset. **B** The proportion of profiles yielding a valid DLMO estimate at each threshold, with failures occurring when all melatonin values remained either below or above the threshold.

The number of profiles yielding a valid DLMO estimate varied by threshold value (see Tbl. 8 and Tbl. 9). In both datasets, failed estimates occurred when melatonin concentrations remained entirely sub- or supra-threshold. All 24 profiles in the Heinrichs and Spitschan (2025) dataset produced valid estimates at thresholds up to 5 pg/mL, with two profiles excluded at 10 pg/mL. In contrast, the Blume et al. (2024) dataset showed success rates ranging from 76% at 2 pg/mL to ≥95% at thresholds of 3 pg/mL and above.

Note that all ΔDLMO values in Fig. 9 are expressed relative to the DLMO at 2 pg/mL, so the maximum number of comparable profiles in Appendix H.2, Tbl. 10 and Tbl. 11 is capped by the number of profiles with a valid estimate at the 2 pg/mL threshold. As a result the *n*_compared_ values in those tables are lower than the corresponding counts *n* in the success tables (Tbl. 8 and Tbl. 9).

To assess the robustness of DLMO estimates to missing or unreliable data, we performed a series of perturbation analyses designed to simulate common deviations from ideal sampling conditions. These included (i) single-point deletions, (ii) multiple-point deletions, and (iii) the addition of noise to either the melatonin concentration, the sampling time, or both. Together, these scenarios reflect challenges frequently encountered in real-world data collection, such as forgotten or missed samples, delays in experimental protocols, and intra-assay variability.

#### Single-point deletions

##### Rationale

Missed or invalid samples are not uncommon in melatonin sampling protocols and may affect the stability of DLMO estimation. To assess how sensitive estimates are to localized data loss, we evaluated the effect of systematically deleting individual samples.

##### Methods

For each melatonin profile, a baseline DLMO was estimated from the complete time series. Then, each data point was systematically deleted, one at a time, and the DLMO was re-estimated for each iteration. The resulting shift in DLMO (perturbed minus baseline) was recorded and associated with the timing of the deleted sample, expressed relative to the baseline DLMO. These relative deletion times were then grouped into 15-minute bins, and the mean and standard deviation of the DLMO shift within each bin were computed across all profiles. Results were visualized using a ribbon plot and heatmap to highlight zones of stability and sensitivity in the melatonin curve. Although deletion effects were computed across the full extent of each profile (e.g., from −1065 to +570 minutes relative to baseline DLMO in Heinrichs and Spitschan (2025) and −375 to 345 minutes in Blume et al. (2024)), we limited visualizations to the [− 120, +120] minute range for visual clarity, as deviations outside this window were negligible.

##### Results

Aggregated across profiles for each dataset, the analysis revealed a critical window of heightened sensitivity within approximately *±*30 minutes of the estimated DLMO, where deleting a single timepoint produced the largest shifts in estimated timing (see Fig. 10). In contrast, deletions during the stable base or late-rise phases had minimal effect. In the Heinrichs and Spitschan (2025) dataset, DLMO estimates were most sensitive to deletions within a window spanning approximately −15 to +60 minutes relative to the baseline DLMO estimate. The largest mean deviation occurred at +15 minutes (−0.6161 *±* 0.7720 hours), followed closely by deletions centered around the baseline DLMO estimate (0 minutes bin: −0.5706 *±* 0.8094 hours) and at −15 minutes (−0.5265 *±*0.8724 hours). Deletions further from DLMO, such as in the baseline phase (− 120 to −60 minutes), had negligible influence, with mean changes typically under *±*0.030 hours and SD below 0.130 hours.

**Figure 10.**
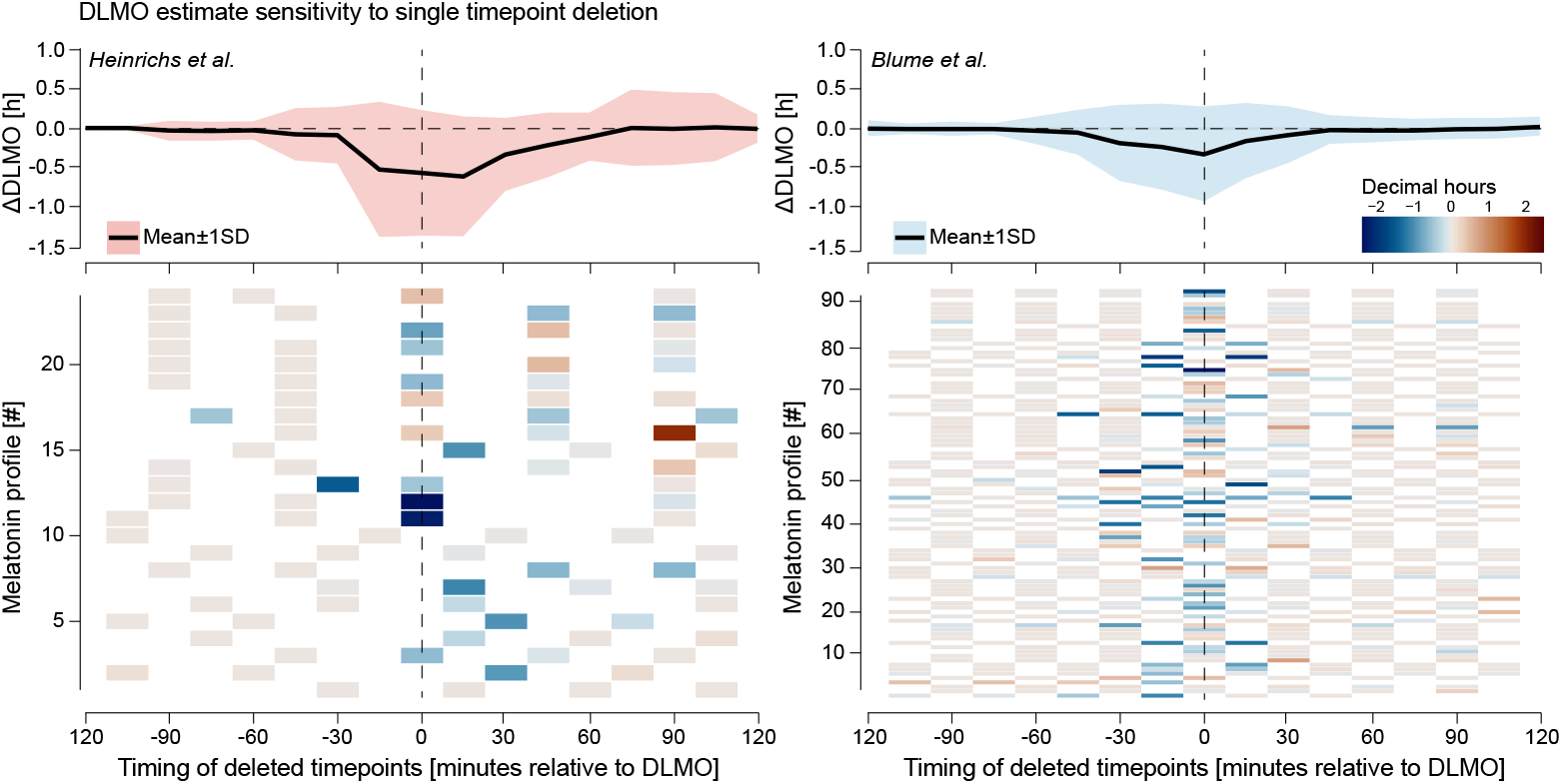
Sensitivity of DLMO estimates to single timepoint deletion for the Heinrichs and Spitschan (2025) (**left**, red) and Blume et al. (2024) (**right**, blue) datasets. For each profile, the DLMO shift resulting from deleting each timepoint was computed relative to the baseline DLMO, with deletion times expressed in minutes from that baseline. **Top panels** Mean (black line) ±1 SD (shaded area) of these shifts across all profiles, binned in 15-minute intervals. **Bottom panels** Per-profile heatmaps of the DLMO shift, with warm colors indicating later and cool colors earlier estimates. In both datasets, a narrow window within approximately *±*30 minutes of DLMO exhibited the greatest sensitivity to deletion, while timepoints further away had little influence.

Sensitivity declined again beyond +60 minutes, though moderate variability persisted (SD = 0.180–0.490 hours).

In the Blume et al. (2024) dataset, sensitivity peaked slightly earlier, with the greatest mean deviation at 0 minutes (−0.3323*±*0.6135 hours), followed by −15 minutes (−0.2380*±*0.5563 hours) and −30 minutes (−0.1901*±*0.4930 hours). Deletions before −60 minutes or after +60 minutes had little impact, with mean shifts below *±*0.030 hours and SD consistently under 0.140 hours (e.g., −120 minutes: mean =− 0.0003 *±* 0.1078 hours; +120 minutes: mean = +0.0249 *±* 0.1227 hours).

The ribbon plots and heatmaps in Fig. 10 illustrate these dynamics across individual profiles, revealing how a small subset of samples, especially those near the DLMO inflection point, can disproportionately affect the final estimate.

#### Multi-point deletions

##### Rationale

Multiple missed samples are a common occurrence in melatonin studies, arising from protocol deviations, participant noncompliance, or technical assay failures. To assess the impact of more extensive data loss, we simulated scenarios in which multiple samples were deleted.

##### Methods

For each melatonin profile, a baseline DLMO was estimated from the complete time series. Perturbation replicates were then generated by randomly deleting 10%, 20%, 30%, 40%, or 50% of the original samples. Each deletion level was repeated 20 times per profile to account for variability in deletion patterns (see Appendix I, Fig. 16 for example profile deletions). DLMO was re-estimated for each replicate, and the deviation from the baseline estimate (ΔDLMO) was recorded.

##### Results

As the proportion of deleted samples increased, both datasets showed progressively larger deviations in estimated DLMO and greater variability across replicates (see Fig. 11 and Appendix H.3 for Tbl. 12 and Tbl. 13). In the Heinrichs and Spitschan (2025) dataset, the average DLMO shift grew from −0.0409 *±* 0.4094 hours) at 10% deletion to −0.3022 *±* 1.1233 hours) at 50% deletion. Estimates remained stable at lower deletion levels, with *>*99% of replicates yielding valid DLMO estimates up to 40% deletion. However, at 50% deletion, the success rate dropped slightly to 97.1%, indicating occasional loss of DLMO detectability.

**Figure 11.**
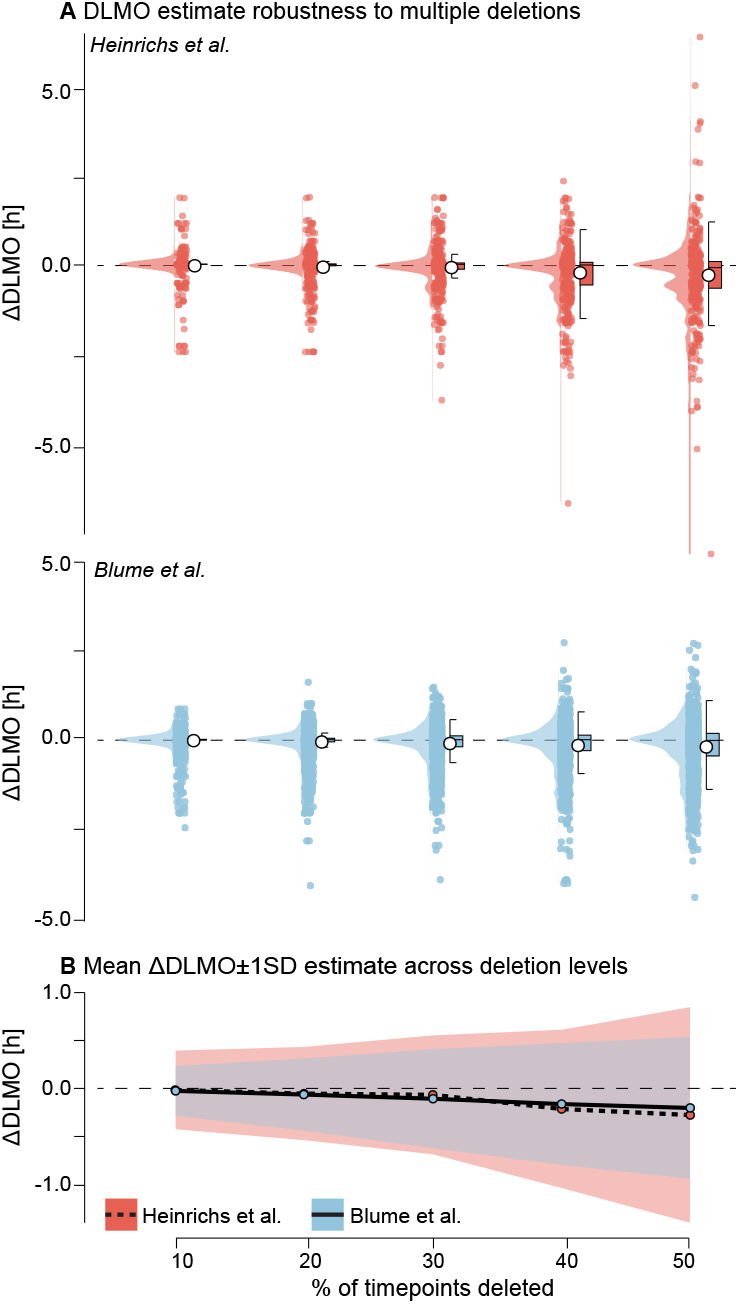
Robustness of DLMO estimates to multiple timepoint deletions for the Heinrichs and Spitschan (2025) (red) and Blume et al. (2024) (blue) datasets. **A** Distribution of ΔDLMO values (difference from baseline) across 20 replicate deletions at each deletion level, with points representing individual replicates, boxplots indicating median and interquartile range, and violins showing the distribution. **B** Mean ΔDLMO (line) *±* 1 SD (shading) across deletion levels for each dataset. In both datasets, greater proportions of deleted samples led to larger deviations and greater variability, with stability at low deletion levels and marked increases above 40% deletion.

In the Blume et al. (2024) dataset, DLMO estimates were initially more stable, with mean shifts of −0.0264 hours *±* 0.2612 hours) at 10% deletion. But sensitivity increased at higher deletion levels, reaching an average deviation of −0.2045 *±* 0.7533 hours) at 50% deletion. Here, the drop in estimation success was more pronounced, decreasing from 95.6% at 10% deletion to 90.8% at 50%. In both datasets, deletions that prevented a DLMO estimate occurred when all remaining timepoints after deletion were either sub-threshold or supra-threshold, precluding a threshold crossing required by the hockey-stick algorithm.

Across profiles, the shift in estimated DLMO increased gradually with deletion percentage, with variability widening as more samples were removed. At 40% data loss, average deviations exceeded 30 minutes, indicating substantial instability. These patterns are visualized in violin plots, which show the full distribution of ΔDLMO across replicates, and summary plots showing the mean DLMO shift with standard deviation across replicates (see Fig. 11).

#### Noise perturbations

##### Rationale

Experimental data are rarely noise-free: imprecise sampling times, participant nonadherence, and intra-assay measurement error can all perturb melatonin profiles. Such noise may affect the stability of DLMO estimation, yet its tolerance bounds remain poorly defined. To quantify robustness under these conditions, we introduced controlled perturbations along both the temporal and concentration axes of the melatonin signal.

##### Methods

Each melatonin profile was processed under six conditions: (i) a clean, noise-free baseline; (ii–iv) temporal jitter applied by adding Gaussian noise to sampling times, with standard deviations of 5, 10, or 20 minutes; (v) melatonin-only noise was modeled as multiplicative Gaussian noise with a coefficient of variation (CV) of 7.9%, corresponding to the intra-assay precision of the RK-DSM2 radioimmunoassay (NovoLytiX GmbH, Switzerland), such that the standard deviation at each timepoint scaled with melatonin concentration; (vi) combined melatonin noise (CV = 7.9%) and 10-minute temporal jitter. For each condition, 20 replicate profiles were generated by perturbing the original time series and interpolating values using monotonic cubic Hermite interpolation (PCHIP) to preserve signal shape. Negative values introduced by interpolation or noise were deemed physiologically implausible and were clipped to zero, with the number of occurrences recorded. DLMO was re-estimated for each replicate, and deviations from the clean baseline estimate were recorded as ΔDLMO.

##### Results

Across both datasets, noise perturbations introduced modest shifts in estimated DLMO, with deviations increasing systematically with noise magnitude and type (see Fig. 12 and Appendix H.4 for Tbl. 14, and Tbl. 15). The mean absolute DLMO deviation remained below 0.0500 hours for all perturbation types in the Heinrichs and Spitschan (2025) dataset and below 0.1250 hours in the Blume et al. (2024) dataset. Variability, however, increased notably with noise level. For instance, in the Heinrichs and Spitschan (2025) data, standard deviation increased from 0.2137 hours under 5-minute jitter to 0.4940 hours under 20-minute jitter. A similar trend was observed in the Blume et al. (2024) dataset, where variability rose from 0.2745 hours to 0.5155 hours across the same conditions.

**Figure 12.**
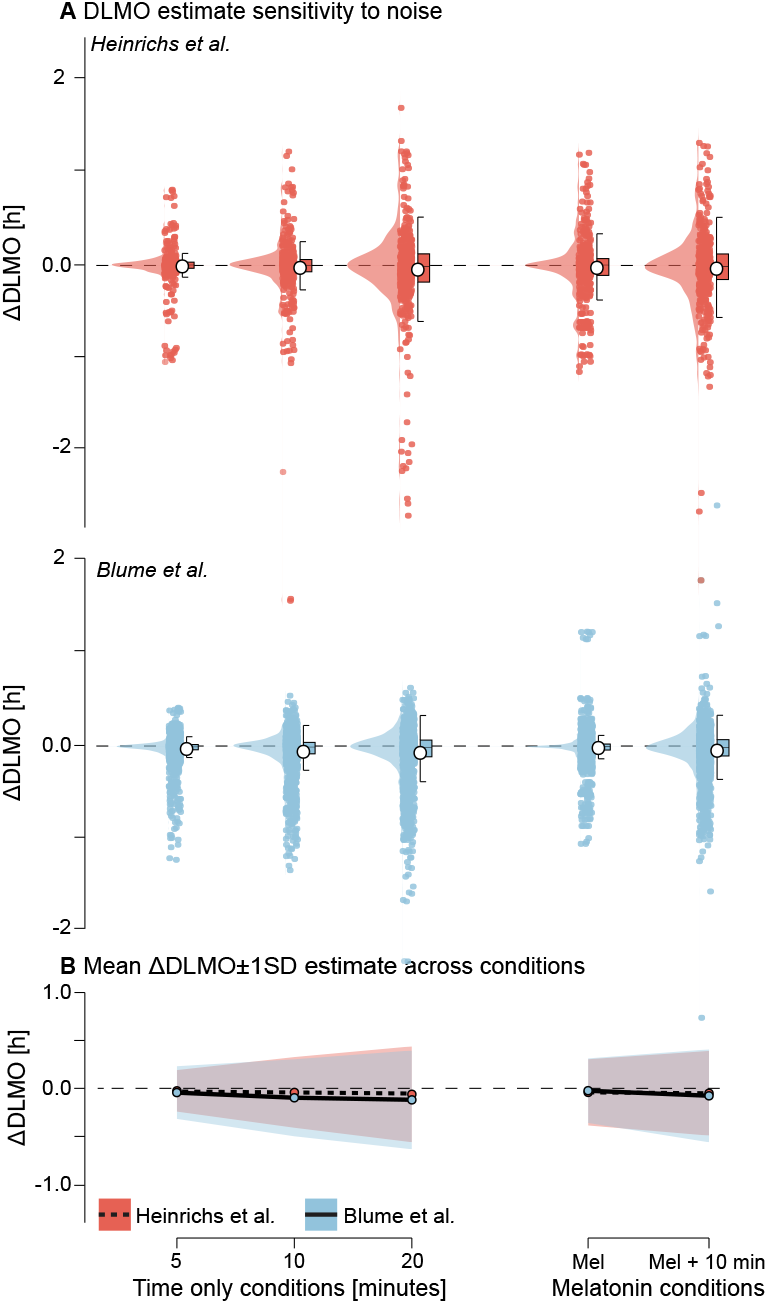
Sensitivity of DLMO estimates to noise perturbations for the Heinrichs and Spitschan (2025) (red) and Blume et al. (2024) (blue) datasets. A Distribution of ΔDLMO values (difference from the noise-free baseline) for each perturbation condition: temporal jitter applied as Gaussian noise to sampling times with different standard deviations; melatonin-only noise applied as multiplicative Gaussian noise with a coefficient of variation (CV) of 7.9% of the concentration; and combined melatonin + 10-minute jitter noise. Points represent individual replicates, boxplots indicate median and interquartile range, and violins show the distribution across 20 replicates per profile. B Mean ΔDLMO (line) ±1 SD (shading) for each noise condition. Across both datasets, DLMO estimates remained largely stable, with variability increasing systematically with noise magnitude and timing jitter having greater impact than melatonin-only noise. Occasional failures to produce a valid DLMO estimate in the Blume et al. (2024) occurred when all perturbed melatonin values were above or below threshold

**Figure 13.**
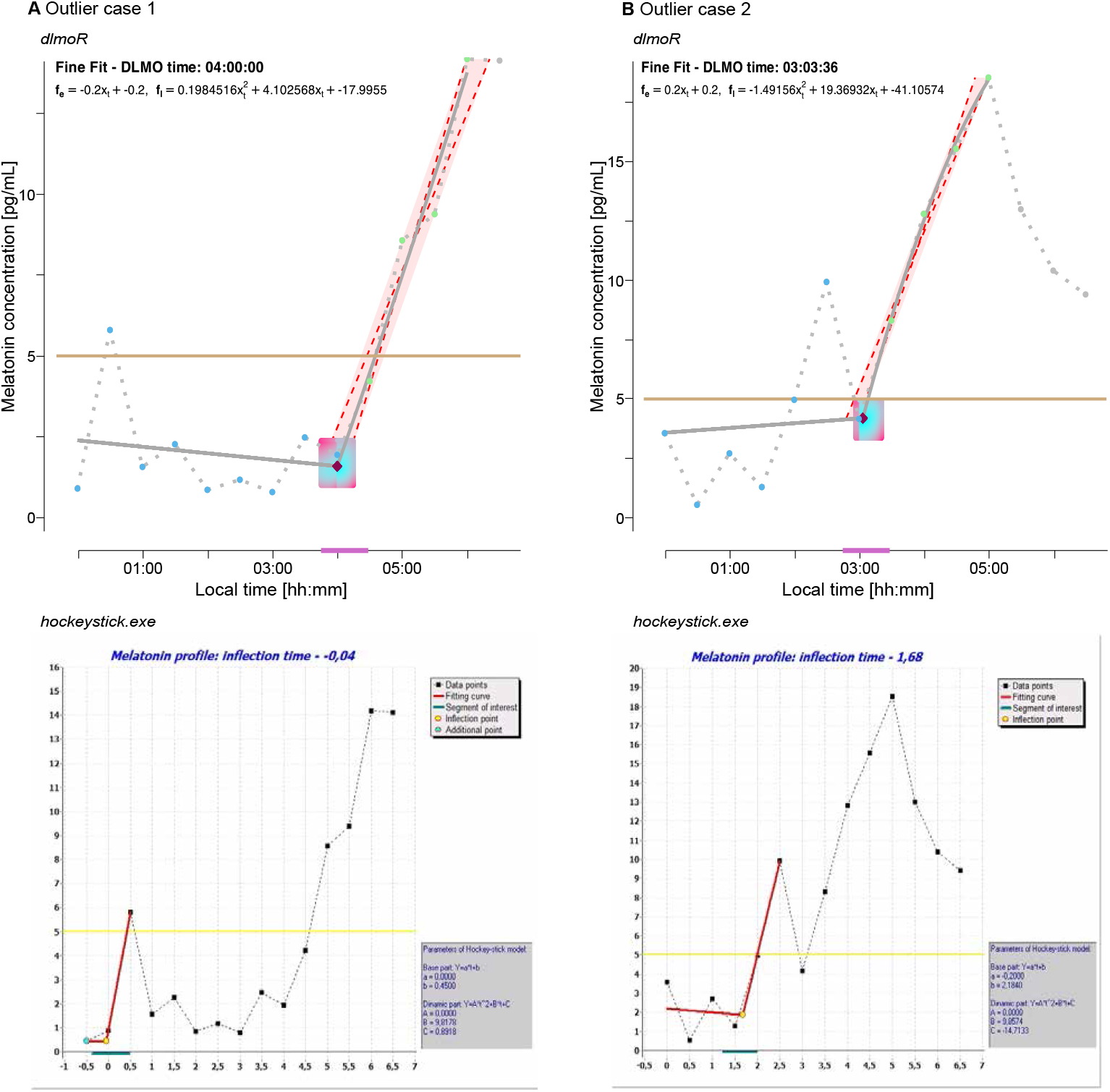
The only two major outlier cases among 112 profiles, showing large discrepancies between *dlmoR* (top row) and *hockeystickexe* (bottom row) DLMO estimates. **A** Outlier case 1: a single early above-threshold melatonin value was fit by *hockeystickexe*, whereas *dlmoR* fit the later sustained rise, yielding a 80-minute difference in DLMO estimates. **B** Outlier case 2: *hockeystickexe* fit a spurious early rise, while *dlmoR* identified the sustained rise 240 minutes later. **Top panels**: the dark purple point marks the DLMO estimate, solid grey lines the model fits, the ochre horizontal line the threshold, the light purple line the ROI range for the coarse fit, and shaded areas the parallelogram used to trim ascending segments. **Bottom panels**: yellow points mark DLMO estimates, solid red lines the model fits, yellow horizontal line the threshold, and the green line the ROI range for the coarse fit.

**Figure 14.**
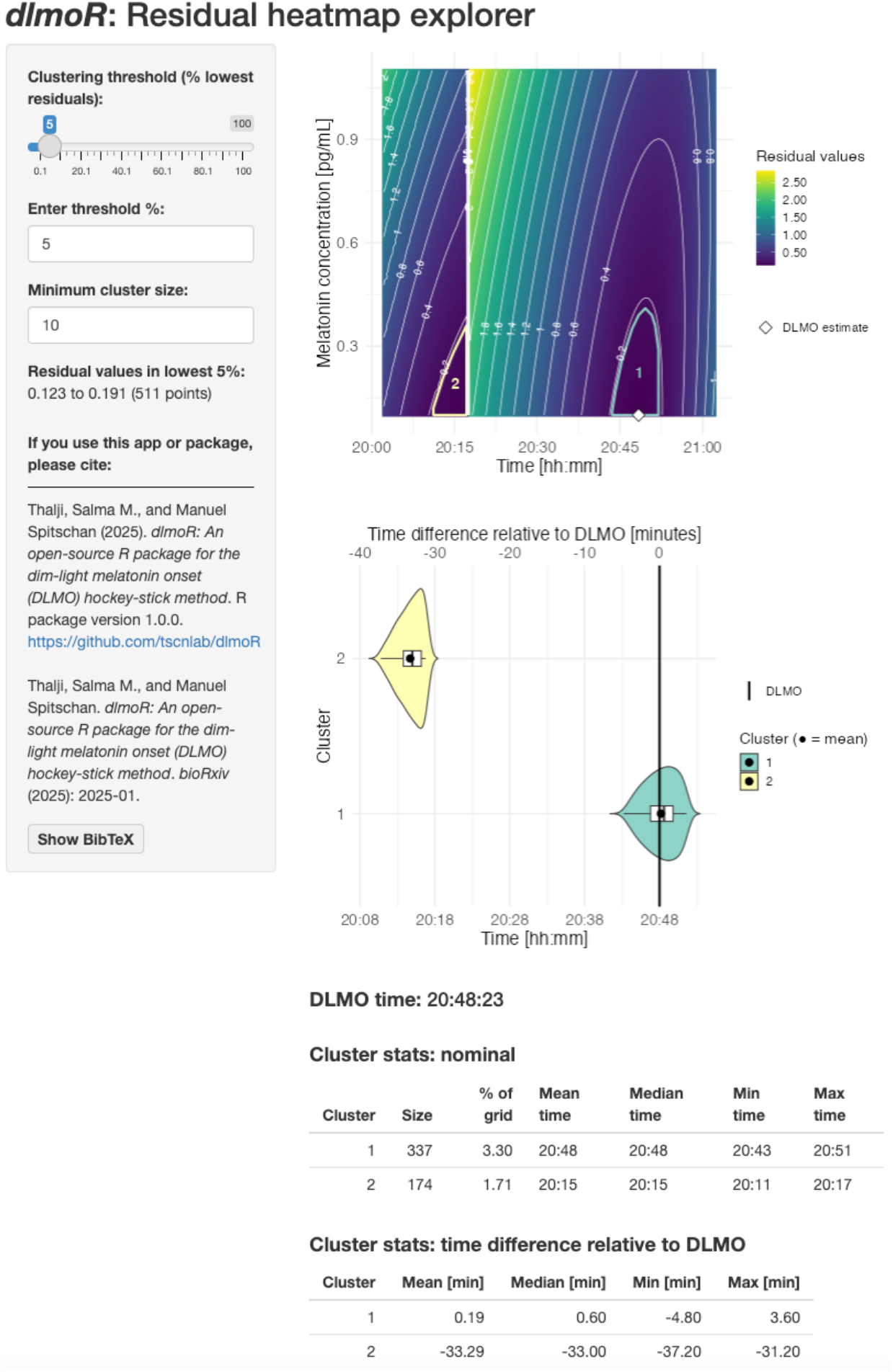
Example output from the *dlmoR: Residual heatmap explorer* Shiny app, a *post-hoc* tool that takes the DLMO data structure output from *dlmoR* as input and visualizes the surrounding residual landscape of the ROI. It allows users to assess whether the landscape is unimodal or multimodal, and how focused or diffuse it is—offering insight into how definitive the DLMO fit may be. **Top left**: The control panel allows users to adjusts the clustering threshold—via sliding bar or numeric input—to set the percentage of lowest residual values included in clusters and the minimum cluster size. **Top right**: The heatmap shows residual values for a single melatonin profile, with contour lines indicating magnitude. Colored regions mark clusters within the specified threshold, and the white diamond marks the DLMO estimate. **Middle right** The violin plot shows candidate DLMO times per cluster, relative to the estimate. **Bottom**: Summary tables list cluster size, proportion of the grid, and timing range.

**Figure 15.**
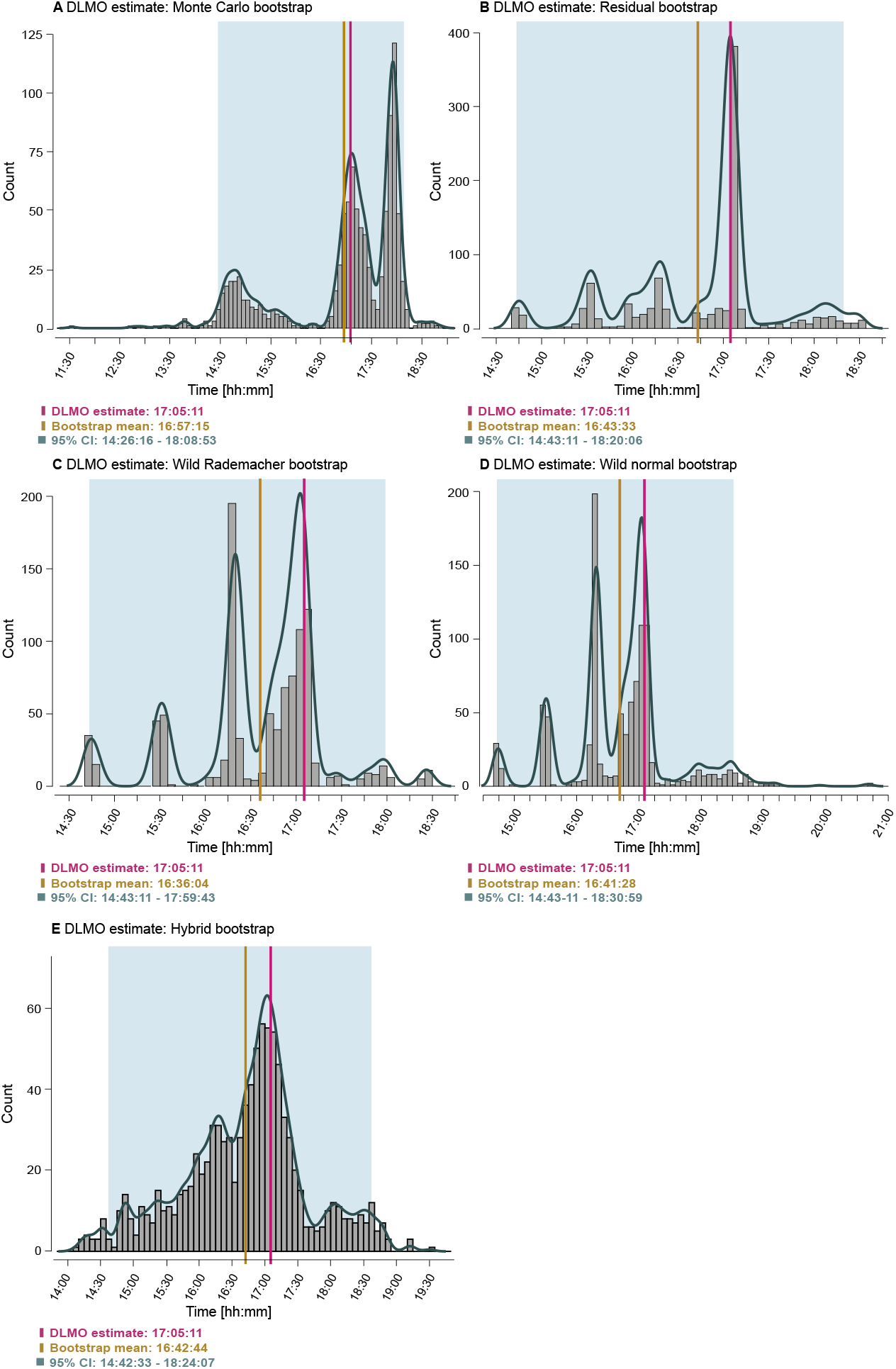
Example distributions of DLMO estimates obtained using five bootstrap methods implemented in dlmoR. Panels show histograms and kernel density estimates for a single melatonin profile under: **A** Monte Carlo bootstrap, which perturbs both sampling times and melatonin concentrations to simulate realistic experimental variability; **B** Residual bootstrap, which resamples model residuals to quantify uncertainty from measurement scatter; **C** Wild bootstrap with Rademacher weights and **D** Wild bootstrap with normal weights, both generalizing the residual approach by applying random weights to residuals; and **E** Hybrid bootstrap, which combines Monte Carlo and residual resampling to capture both timing and residual noise. Vertical magenta lines mark the original DLMO estimate, ochre lines the bootstrap mean, and shaded blue regions the 95% confidence intervals.

**Figure 16.**
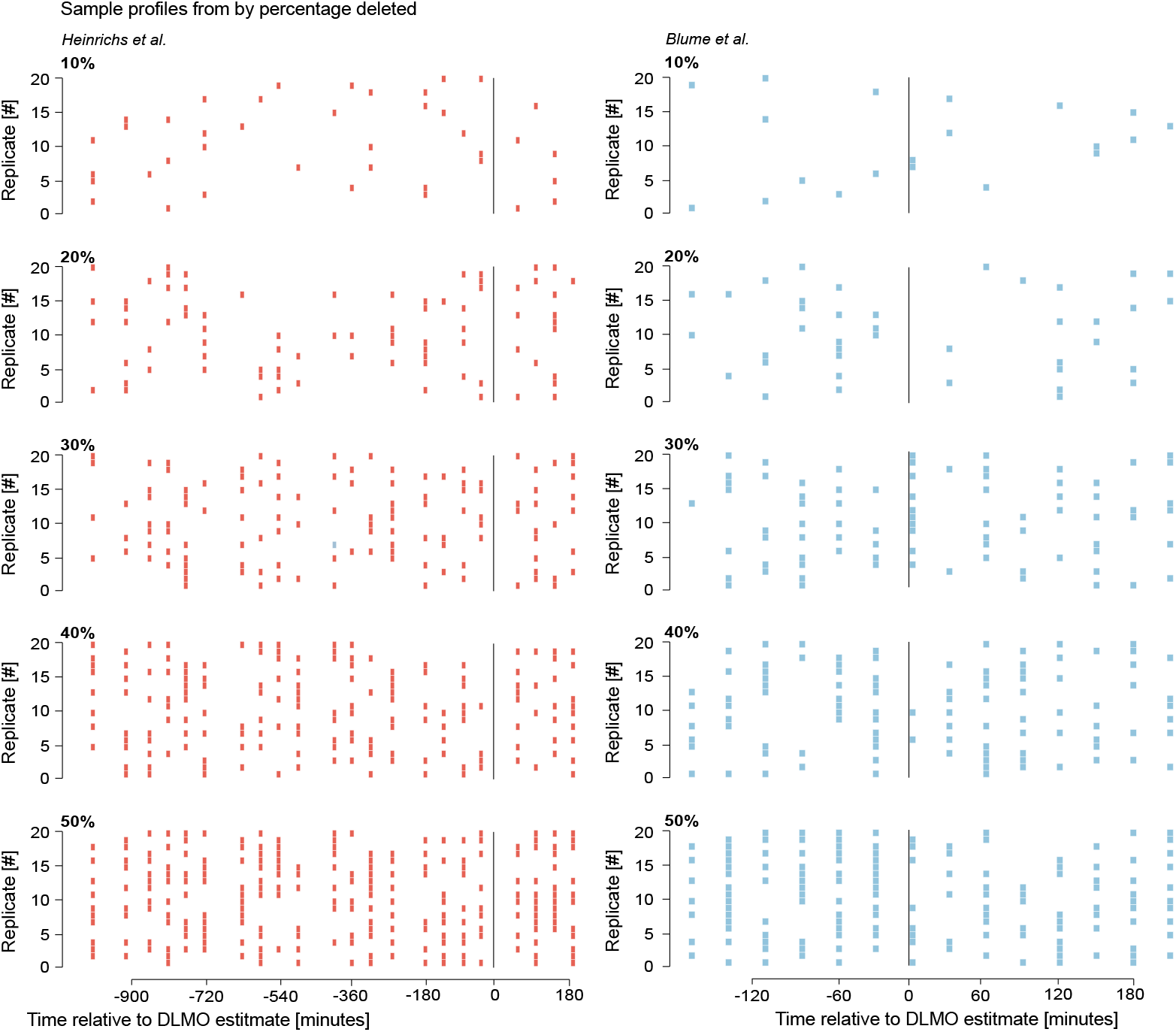
Example raster plots of melatonin profiles after random deletion of 10–50% of samples, for Heinrichs and Spitschan (2025) **(red, left)** and Blume et al. (2024) **(blue, right).** Each row shows one deletion replicate (20 per profile per deletion level), with points marking the remaining samples after deletion, plotted relative to the baseline DLMO estimate from the complete, undeleted profile (vertical black line at time zero). Because deletions were applied at random within each profile, the pattern of missing samples varies between replicates, illustrating how temporal coverage of the melatonin curve is progressively reduced in different ways as the percentage of deleted samples increases. The wider time axis range in the Heinrichs and Spitschan (2025) panel reflects the longer sampling window of those profiles compared to the Blume et al. (2024) dataset. These same deletion replicates form the basis for the multi-point deletion robustness analysis shown in Section *Results: Multi-point deletions*, Fig. 11.

Perturbations that affected only melatonin values (CV = 7.9%) had less impact than timing jitter: melatonin-only noise produced a mean DLMO shift of −0.0303*±*0.3410 hours) in the Heinrichs and Spitschan (2025) dataset, and −0.0273 *±* 0.34 hours) in the Blume et al. (2024) dataset. When both melatonin and timing noise were combined (10-minute jitter + 7.9% CV), variability further increased (SD = 0.4363 for Heinrichs and Spitschan (2025), 0.4849 for Blume et al. (2024)), and the mean shift grew slightly more negative (–0.0396 and –0.0809 hours, respectively).

All 480 runs from the Heinrichs and Spitschan (2025) dataset yielded valid DLMO estimates under all perturbation conditions. In the Blume et al. (2024) dataset, 10 profiles failed to produce estimates in one or more conditions, resulting in fewer runs than the full 1920 possible per condition. As in previous analyses, these failures occurred when melatonin concentrations remained entirely sub-threshold or supra-threshold. Noise perturbations and interpolation introduced negative melatonin values in 216 instances for the Heinrichs and Spitschan (2025) dataset and 88 instances for the Blume et al. (2024) dataset; these were subsequently clipped to zero.

## Discussion

### Concordance between hockeystickexe and dlmoR

The high concordance between *dlmoR* and *hockeystickexe*, supported by high circular correlations and non-significant paired t-tests, demonstrates robust agreement. Descriptive statistics showed small mean differences, with greater variability in the Blume et al. (2024) dataset than in Heinrichs and Spitschan (2025). Time-threshold assessments further indicated that most DLMO estimates were within 15 minutes of each other. The slight tendency for *dlmoR* to produce earlier estimates, also seen in skewness analysis, likely reflects minor numerical differences in the optimization procedure. Because the closed-source *hockeystickexe* uses undisclosed algorithms and initialization parameters, *dlmoR* applies an informed but independently selected optimization method and initial values. In addition, the hockey-stick algorithm derives the DLMO estimate from the single lowest residual within the ROI heatmap; in some analyzed profiles, hundreds of residuals differed by less than 1 *×* 10^−2^ in magnitude. In such cases, even the smallest difference between implementations, such as the exact start and end points of the grid search locations, can change the time point that corresponds the lowest residual, leading to substantial jumps in the estimated DLMO time. These differences are purely technical and do not involve changes to the underlying algorithm structure.

The closed-source hockey-stick implementation Danilenko and Verevkin (2020) has itself been shown to produce DLMO estimates that are, on average, 3.6 *±* 8.0 minutes earlier than expert judgments (see validation on page 5 in Danilenko et al. (2014)). Whether such small offsets are biologically meaningful depends substantially on context and research question: in clinical or chronotherapeutic applications, an hour’s difference could affect treatment timing, and in mechanistic studies with high statistical power, even small shifts may be critical. In contrast, population-level chronobiological research is generally less sensitive to small systematic biases if the

Thalji and Spitschan 15Time only conditions [minutes]Melatonin conditions10520MelMel + 10 min-1.00.01.0B Mean ΔDLMO±1SD estimate across conditions A DLMO estimate sensitivity to noiseHeinrichs et al.-20.02ΔDLMO [h]Blume et al.-20.02ΔDLMO [h]ΔDLMO [h]Heinrichs et al.Blume et al. method is consistent. For perspective, weekday–weekend DLMO differences in adolescents and young adults are typically 30–60 minutes with corresponding changes in sleep timing (Crowley et al. 2024; Zerbini et al. 2021), and naturalistic light exposure can advance DLMO by approximately 2 hours (Wright et al. 2013). The < 5 minute average difference we observed between the *dlmoR* and *hockeystickexe* implementations is therefore likely biologically inconsequential. A definitive evaluation would require a well-defined ground truth, which does not currently exist.

Identified outliers in DLMO estimates between the two methods, underscore the importance of careful result interpretation and the need for users to pre-assess raw data for spurious melatonin patterns before analysis. Beyond replicating the hockeystickexe implementation, dlmoR provides an interactive Shiny-based diagnostic tool for exploring the residual heatmap of the hockey-stick fit and a function for bootstrapped confidence interval estimation. Together, these features facilitate both qualitative and quantitative assessment of DLMO uncertainty and offer insight into the robustness of each estimate.

### Hockey-stick algorithm performance under varying conditions

Our open source implementation of the hockey-stick algorithm enabled us to conduct a series of analyses that probed the internal mechanics of the method by challenging it with controlled changes to sampling, thresholds, data availability, and noise. This approach revealed how specific alterations propagate through the algorithm’s region-of-interest definition and piecewise fit, sometimes producing subtle shifts in DLMO timing and in other cases causing marked deviations or failed estimates. The results set the stage for understanding why certain aspects of study design and data quality exert disproportionate influence on the DLMO outcome.

The resampling analysis showed that the hockey-stick algorithm is sensitive to both overly fine and overly coarse sampling. In the Heinrichs and Spitschan (2025) dataset, variability followed a non-monotonic pattern, with elevated deviations at very short intervals (2-10 minutes) and at coarse intervals (≥ 75 minutes), and minimum variability at intermediate intervals (30-45 minutes). In contrast, the Blume et al. (2024) dataset remained stable up to 30 minutes but showed increasing bias and variability at coarser intervals. This likely reflects differences in the original sampling schedules: the Heinrichs and Spitschan (2025) profiles were collected at intervals close to 45 minutes, so resampling to finer grids amplified the influence of small variations in timing and concentration, whereas the Blume et al. (2024) profiles were sampled every 30 minutes, so undersampling more quickly reduced temporal resolution and potentially masked the true onset time. Oversampling can magnify the influence of noise, while undersampling widens the gaps between points, making the DLMO search less constrained and thus broadening the domain of possible DLMO estimate locations (see threshold analysis below for a detailed discussion of how ROI boundaries affect the search). Our results suggest that intervals in the range of 20-45 minutes may provide an optimal balance between temporal precision and feasibility for this algorithm.

Raising the melatonin threshold from 2 to 10 pg/mL systematically delayed the DLMO and, at higher thresholds, increased variability. This behavior is expected because the hockey-stick algorithm uses the threshold level in two critical ways: (i) it sets the upper bound of the region of interest (ROI), and (ii) it determines the classification of points into *baseline, intermediate*, or *ascending* regions, which in turn defines the temporal limits of the ROI. Changing the threshold therefore alters both the width (i.e. time range) and height (i.e. concentration range) of the search region, as well as the subset of points available for the piecewise-linear fit, thereby directly influencing the estimated break point. The Heinrichs and Spitschan (2025) dataset was notably more resistant to threshold changes than Blume et al. (2024), likely due to the steeper melatonin rise in its profiles. Larger vertical concentration changes between consecutive samples make the baseline–ascending boundary (and thus the ROI limits) less sensitive to small threshold shifts.

Given the strong influence of threshold level on DLMO timing, this choice should be made transparently and, where possible, applied consistently across samples and studies to enable comparability. In most cases, a uniform threshold is recommended; however, in rare instances, individual melatonin profiles may require an alternative, for example, when concentrations never approach the standard value or when unusually high baseline levels obscure the onset. Our recommendations for choosing and reporting a standard threshold level, and handling of exceptional cases, are provided in Appendix J.

The single-point deletion analysis revealed a narrow window of heightened sensitivity spanning just before to shortly after the onset of melatonin rise. Within this range, removing a single sample could shift the estimate by 30–40 minutes or more. This behavior reflects the algorithm’s dependence on ROI boundaries: the ROI begins at the last *baseline* point, includes any *intermediate* points, and extends into the *ascending* phase. Deleting a point that marks one of these transition bounds alters the ROI limits, which in turn redefines the grid search domain and the set of candidate DLMO estimates. By contrast, deleting points well outside this transition zone affects the piecewise-linear fits for individual segments but leaves the ROI bounds intact, thereby limiting their influence on the final DLMO estimate. Multi-point deletions compounded these effects. While removing 10–20% of samples produced limited shifts in the DLMO estimate, deletions exceeding 30% caused substantial instability, with mean deviations exceeding 30 minutes at 40% loss. Deletions affect the estimate both by widening the ROI (due to greater gaps between the remaining points) and by reducing the number of points available to fit each segment, making the break point more susceptible to shifts in position. Profiles with shallow melatonin rises or low peak concentrations were especially vulnerable, and in some cases failed to yield a valid estimate when the profile became entirely sub- or supra-threshold.

Perturbations mimicking temporal jitter and assay noise revealed that the hockey-stick algorithm is particularly susceptible to changes in sample timing. This is because assignment of region labels to points, and thus ROI bounds, depend on sample times and slopes. Small timing shifts can change the classification of points from *baseline* to *intermediate* or even *ascending* (or vice versa), which moves the ROI start earlier or later and alters the search domain. While melatonin concentration noise alone had a smaller effect, the combination of timing jitter and amplitude noise amplified both bias and variability, and in some cases prevented estimation altogether by pushing all points into sub- or supra-threshold ranges.

This in-depth stress-test analysis of the hockey-stick algorithm demonstrates that its performance is shaped in systematic but uneven ways by analytical choices and common data imperfections. In doing so, it provides a framework for understanding the robustness, limitations, and optimal use of the hockey-stick algorithm across diverse experimental contexts. By deliberately perturbing sampling schedules, threshold levels, data completeness, and signal fidelity, we identified both the conditions under which DLMO estimates remain stable and the scenarios that introduce substantial bias or prevent estimation altogether. Its reliance on a fixed threshold, explicit ROI bounds, and a piecewise-linear fit makes this algorithm sensitive to sampling frequency, threshold selection, data loss, and sampling error. To mitigate these vulnerabilities, we recommend that researchers intending to use the hockey-stick algorithm for DLMO estimation: (i) select sampling intervals that capture the rising phase with sufficient resolution; (ii) pre-specify and justify the threshold concentration; (iii) minimize missing data in the *±*1-hour window around the expected DLMO; and (iv) apply quality checks to flag profiles at risk of failed estimation and, where necessary, adjust parameters accordingly. Implementing such safeguards can improve reproducibility and ensure that DLMO estimates reflect accurate onset times within realistic data quality constraints.

### Future directions in DLMO estimation

The development of the hockey-stick algorithm by Danilenko et al. (2014) was a key step forward in providing an objective framework for determining DLMO, addressing the subjectivity inherent in earlier approaches. However, upon closer examination, this method is not as definitively objective as it might initially appear. The sensitivity of its optimization process, particularly to initialization and local minima, reveals underlying challenges, especially in complex tasks like the multi-constrained optimization of the parallelogram used to select the subset of ascending-phase points for fitting with the hockey-stick model. This sensitivity, coupled with the diffuse nature of the ROI heatmap, highlights that the solutions derived are not singular or definitive but instead represent a range of plausible possibilities. Rather than masking this inherent uncertainty, we propose a shift toward approaches that maintain objectivity while transparently representing the spread of potential solutions. The diffuse heatmaps often found in our profile analyses suggest that the DLMO is not a precise timestamp but rather a region of highest likelihood.

One promising refinement within the same general class of piecewise-linear models is the use of the *segmented* package in R (Rne 2008), which implements the approach described by Muggeo (2003) for estimating breakpoints and their uncertainty in regression models. Unlike the discrete search grids or fixed candidate points used in the hockey-stick algorithm, *segmented* treats breakpoints as model parameters, iteratively estimating them alongside regression slopes using a score-based method (Fasola et al. 2018). This framework supports formal statistical inference, including standard errors and confidence intervals for breakpoints, with more recent advances improving interval estimation through smoothed score-based approaches (Muggeo 2017). By allowing multiple breakpoints, varying slopes before and after the onset, and hypothesis testing to evaluate the significance of added breakpoints, *segmented* addresses a key limitation of the traditional hockey-stick algorithm: the lack of formal statistical inference and uncertainty quantification for the estimated onset point. While still grounded in the conceptual simplicity of piecewise-linear transitions, *segmented* offers a more rigorous framework for breakpoint estimation, making it better suited for datasets where the rise in melatonin is gradual or noisy.

Despite these advantages, *segmented*, like other breakpoint-based methods, ultimately produces a single best estimate of the onset point, supplemented by a confidence interval. This still frames DLMO as a fixed timestamp rather than a distribution over possible values. A fully probabilistic approach, such as a Gaussian process model (GPM) (Rasmussen and Williams 2006), could better capture this uncertainty, modeling DLMO as a probabilistic distribution over a plausible range rather than a single optimal point. GPMs offer several advantages in this context. By representing melatonin profiles as smooth, non-linear functions, they provide flexibility and adapt directly to the data without imposing rigid assumptions like breakpoints. They also naturally account for uncertainty by providing credible intervals for potential DLMO regions, offering a more transparent and nuanced view of melatonin onset. Additionally, GPMs avoid the pitfalls of optimization-based methods, such as sensitivity to initialization or local minima, by calculating distributions over plausible solutions, making them particularly well-suited for noisy or complex datasets.

Alternatively, generalized additive mixed models (GAMMs) (Pedersen et al. 2019) provide another flexible, data-driven approach for estimating DLMO. Unlike Gaussian process models, which take a fully probabilistic approach, GAMMs model melatonin changes as smooth, continuous curves that adapt to the underlying patterns in the data while incorporating fixed and random effects. GAMMs are particularly well-suited to capturing gradual or complex trends without relying on abrupt transitions, such as breakpoints in piecewise models. They also account for individual variability through random effects, allowing the model to adjust for differences across individuals, such as chronotype variation, while still identifying shared patterns. Additionally, GAMMs can pinpoint the sharpest increase in melatonin levels using derivatives of the smooth function, providing a precise and objective estimate of DLMO without relying on predefined thresholds.

As we continue to develop the *dlmoR* package, we aim to build it into a modular suite of DLMO estimation methods, enabling users to select, apply, and directly compare multiple approaches within a single framework. Such flexibility would support both methodological transparency and the exploration of how different modeling assumptions influence DLMO estimates.

Equally important to expanding methodological breadth is ensuring the reproducibility and accessibility of these tools. Reproducibility is a cornerstone of scientific research, enabling findings to be validated and extended by others (Harrison et al. 2021). Yet many scientific software tools lack the transparency, accessibility, and adaptability necessary to meet these standards, hindering broader adoption and integration into diverse research workflows (Wilson et al. 2017; Harrison et al. 2021). Our open-source R package addresses these challenges by promoting reproducible large-scale analyses and facilitating community-driven contributions. Unlike existing standalone.exe software, which limits modifiability and flexibility, this package adheres to the principles of findability, accessibility, interoperability, and reusability (FAIR), fostering an ecosystem for collaborative development and innovation (Lee et al. 2021; Wilson et al. 2017). Through modular design, clear documentation, version control, and continuous integration, dlmoR is designed for both immediate usability and long-term sustainability. By releasing it openly, we aim to equip researchers with a robust, extensible platform for melatonin analysis and a foundation for future methodological advances.

## Conclusion

In this article, we introduced the *dlmoR* package, an open-source R implementation of the hockey-stick algorithm (Danilenko et al. 2014) licensed under the MIT license. Our implementation reproduces the results of the original closed-source executable while making the algorithm transparent and extensible. Through a series of systematic stress tests, we used *dlmoR* to evaluate the robustness of the hockey-stick algorithm, showing that its performance depends in uneven but interpretable ways on sampling schedules, threshold choice, data completeness, and signal fidelity. These analyses provide practical guidance for use of the hockey-stick method for DLMO estimation across experimental contexts. The *dlmoR* package thus establishes a reproducible and sustainable foundation for melatonin-based circadian phase estimation and a modular platform for incorporating future quantitative approaches to DLMO detection.

## Supporting information

Graphical abstract

## Acknowledgements

We wish to thank Dr. Konstantin (Kostya) Danilenko (1962-2023) (Putilov 2023) for detailed discussions of the algorithm and providing a natural-language description of its implementation.

## Declaration of generative AI and AI-assisted technologies

During the preparation of this report, the authors used ChatGPT 4o to improve the readability and language of the manuscript. The authors reviewed and edited the content as needed and take full responsibility for the content of the publication.

## Appendix A Pseudocode of the hockey-stick algorithm implemented in dlmoR

### Algorithm 1

Define *base* region of the melatonin profile

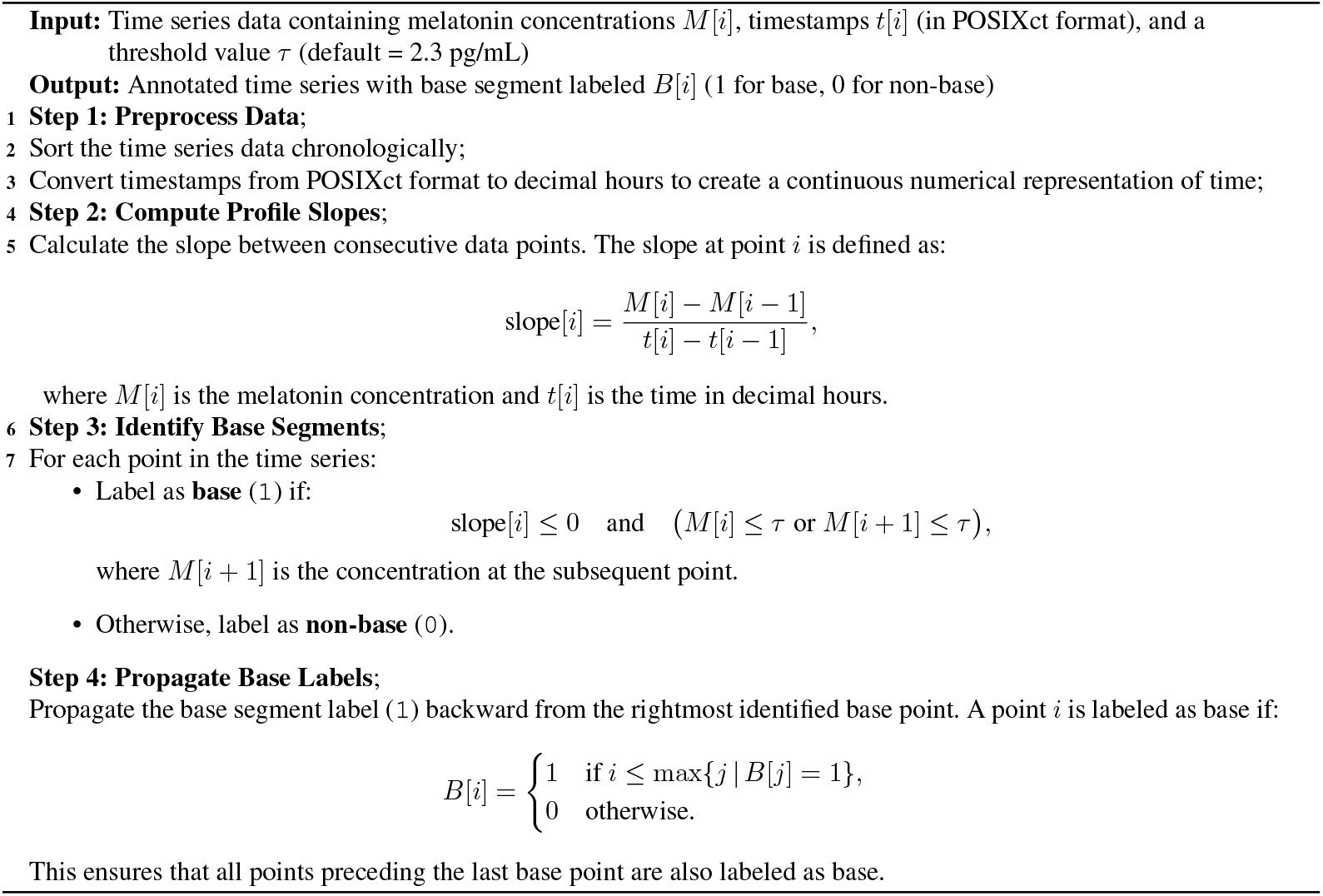

### Algorithm 2

Define *ascending* region of the melatonin profile

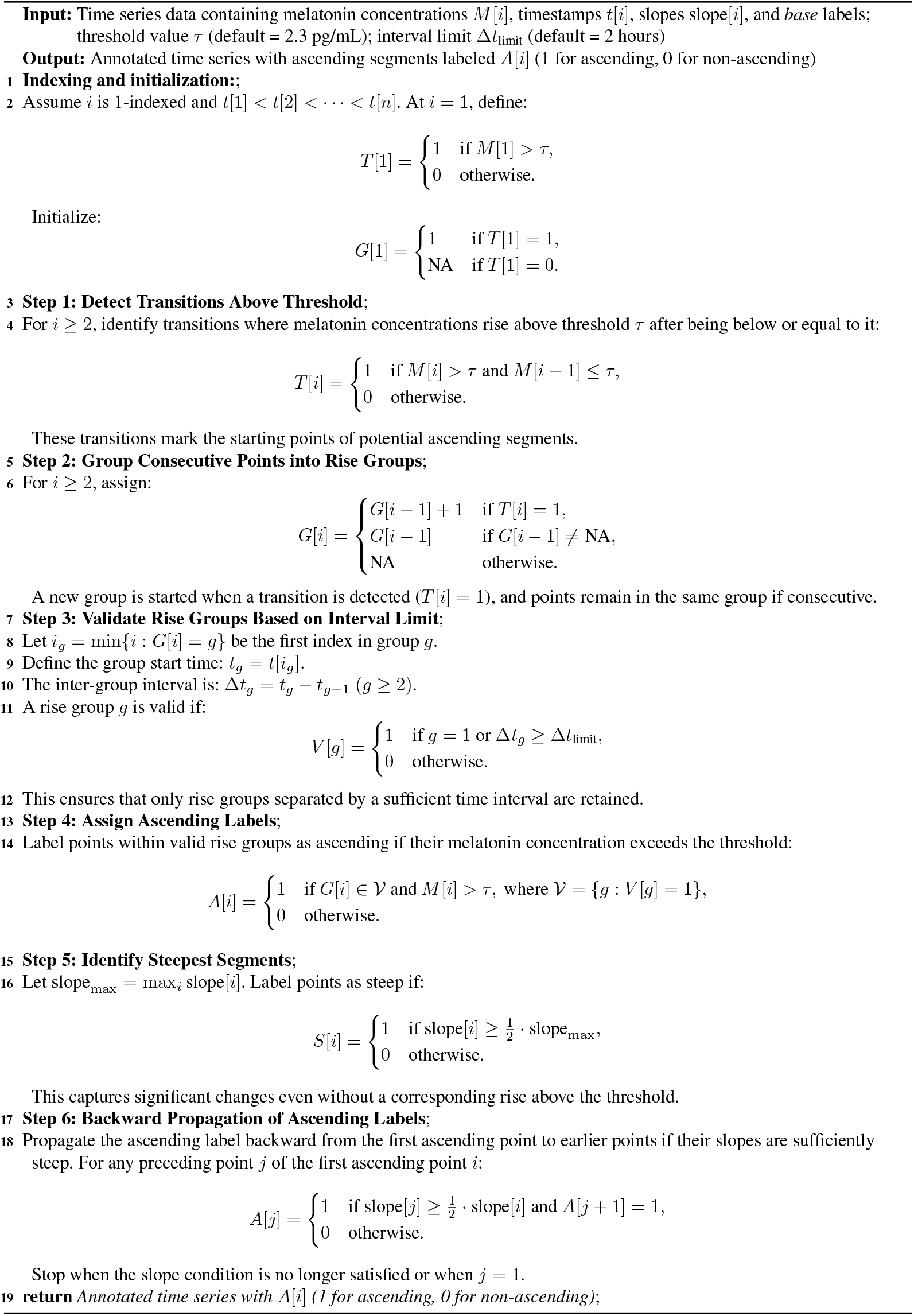

### Algorithm 3

Truncate *ascending* region

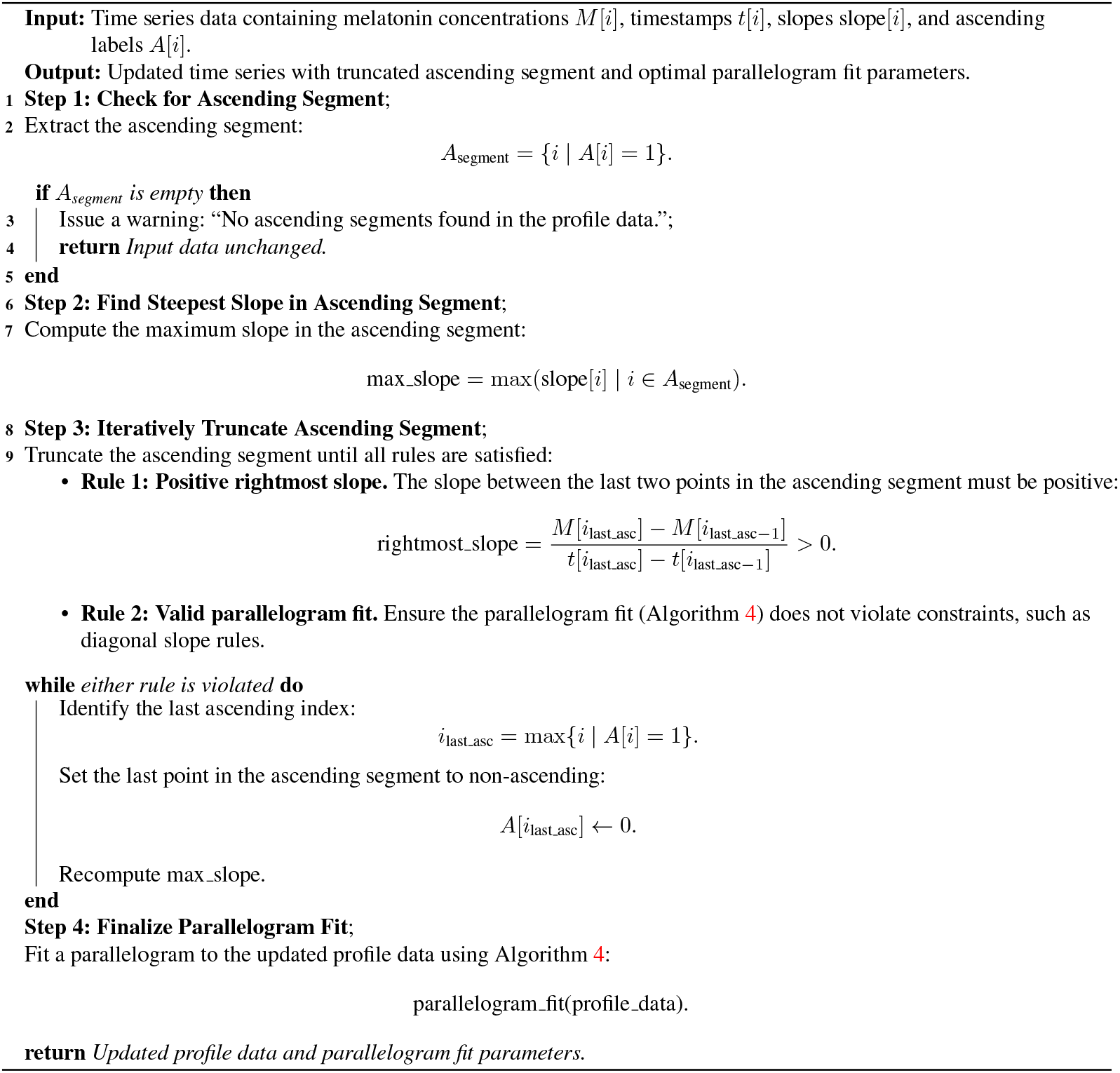

### Algorithm 4

Fit optimal parallelogram to enclose points with minimal area

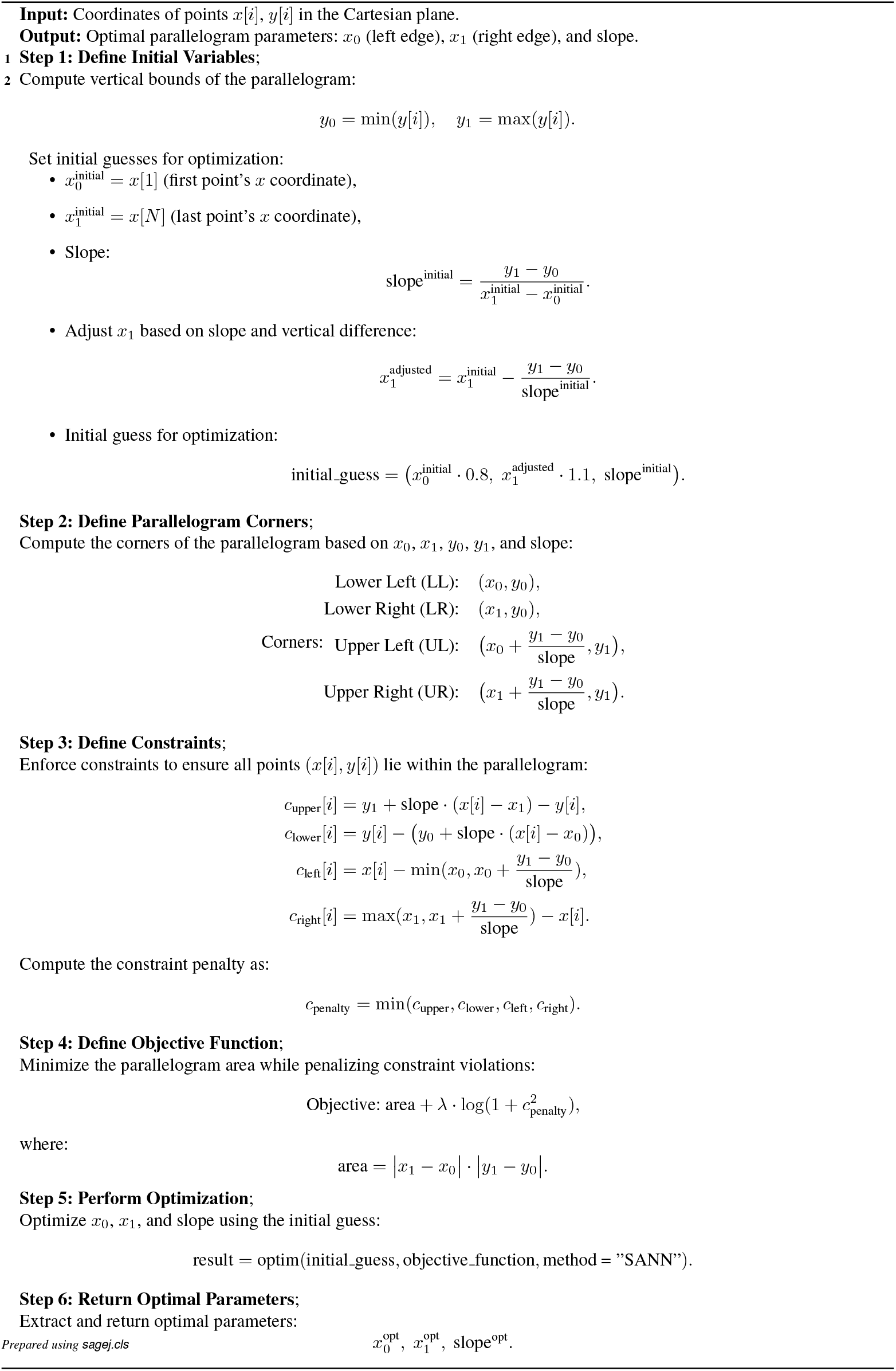

### Algorithm 5

Add a segment to the left of the *ascending* region

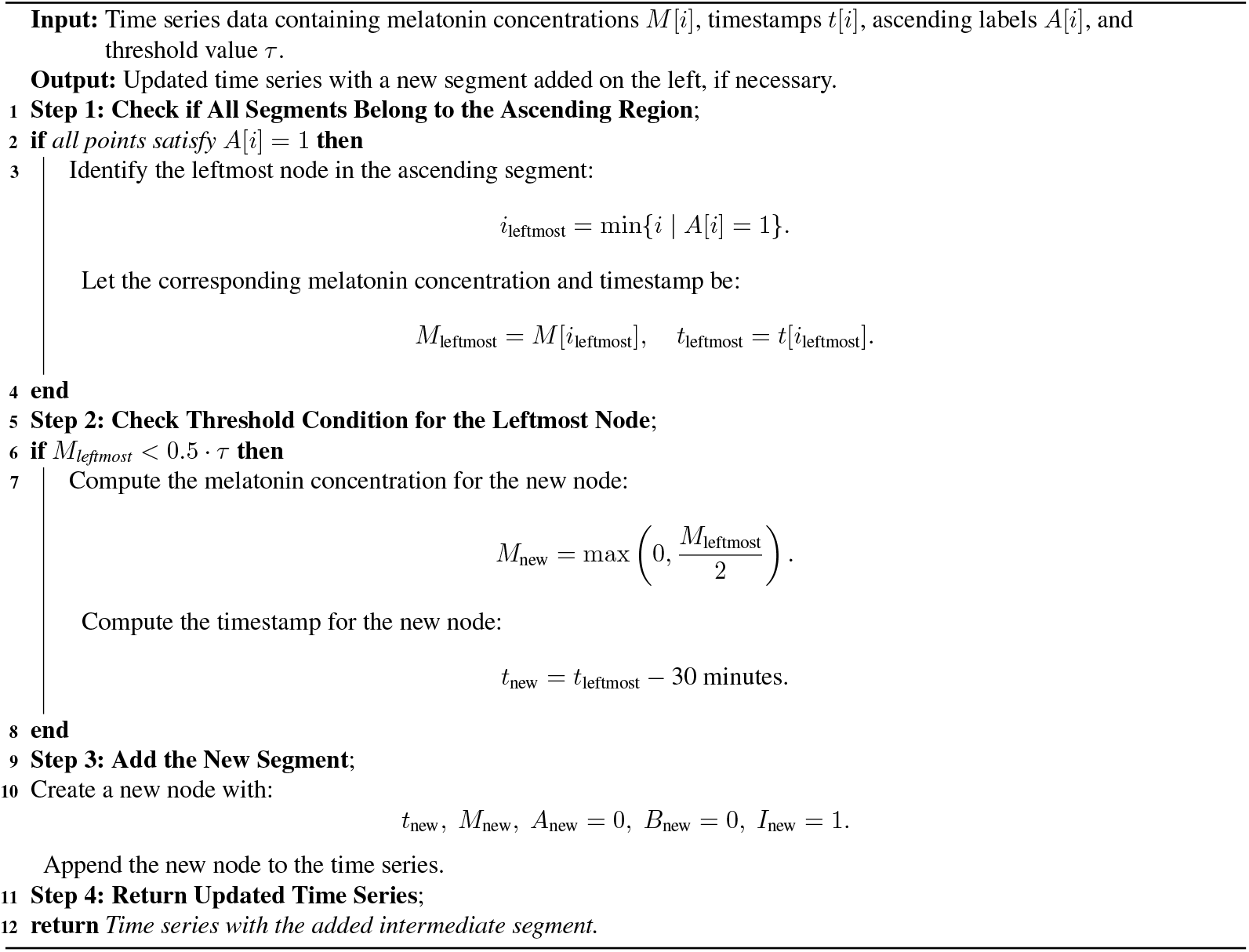

### Algorithm 6

Truncate base region based on threshold

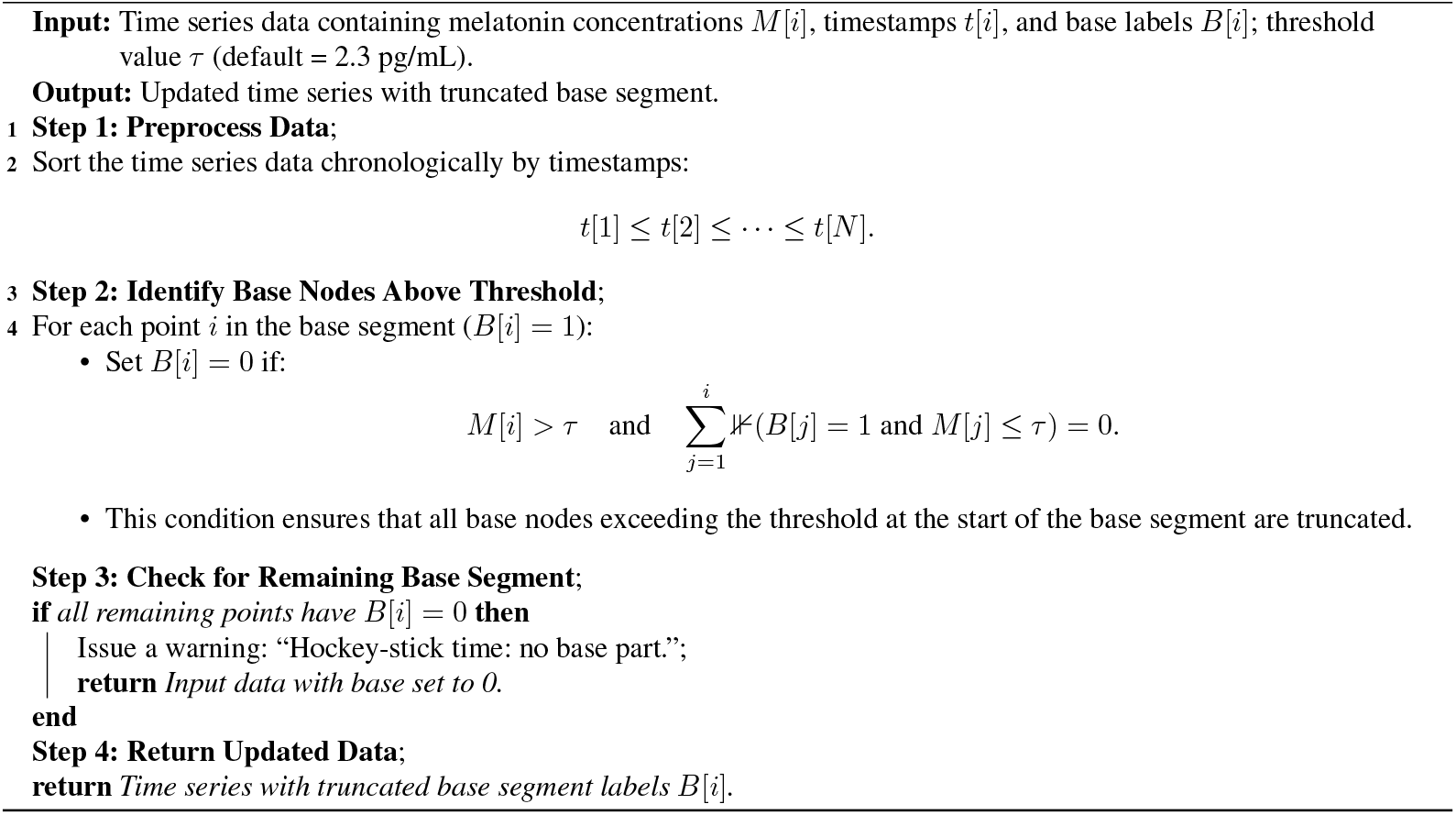

### Algorithm 7

Define *intermediate* segment in a melatonin profile

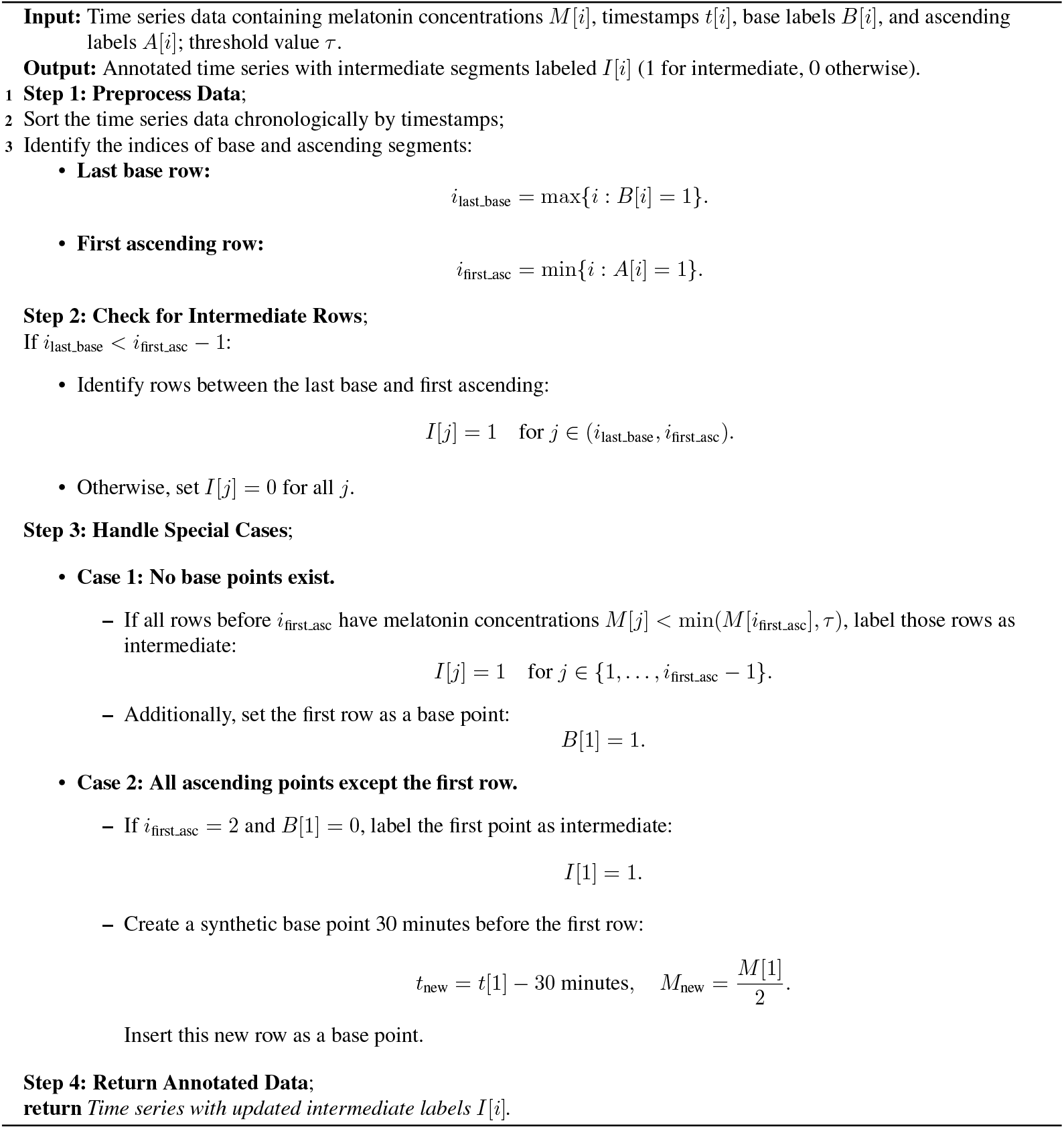

### Algorithm 8

Compute region of interest (ROI) for DLMO point search

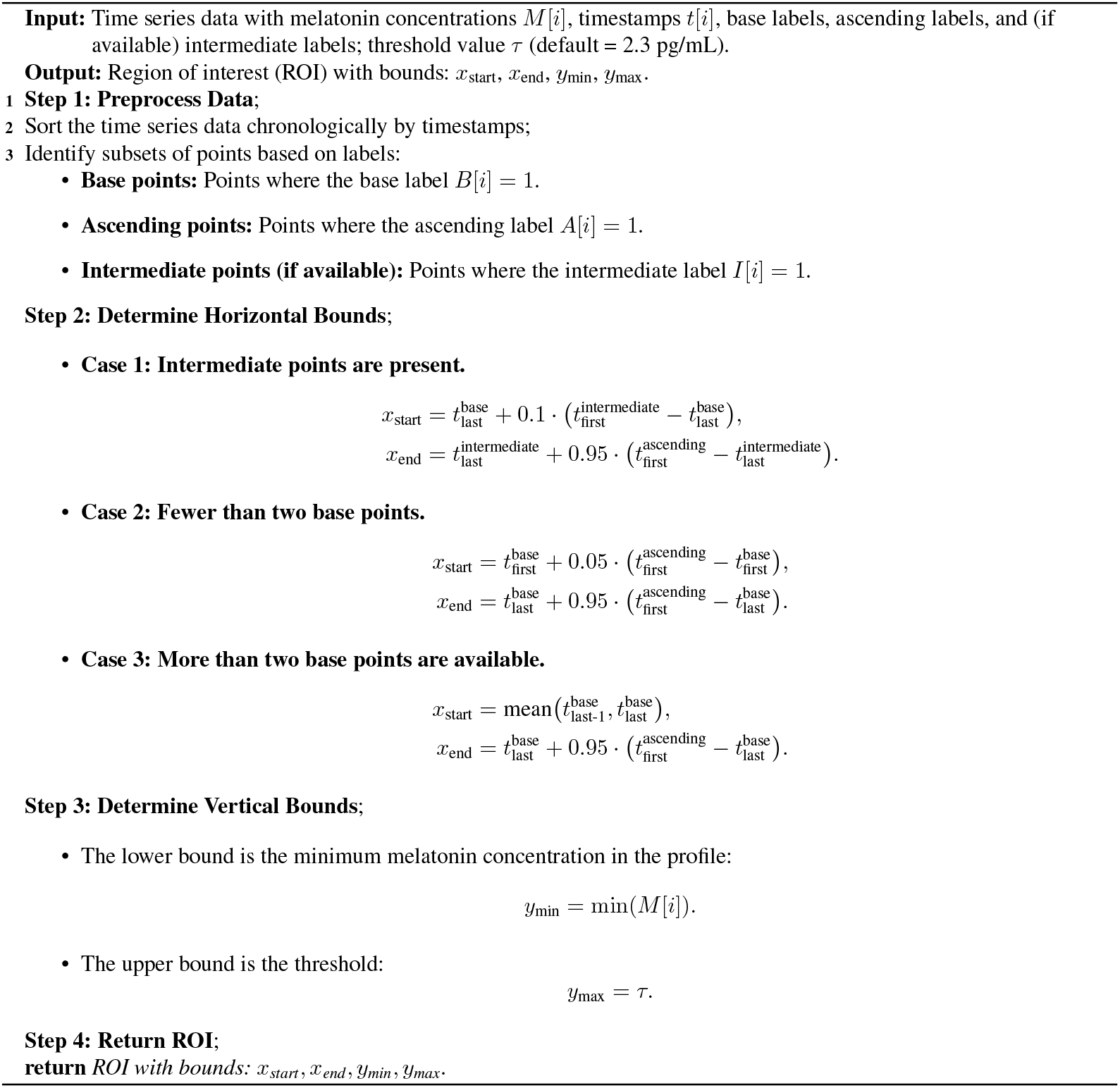

### Algorithm 9

Determine DLMO point using coarse and refined grid search

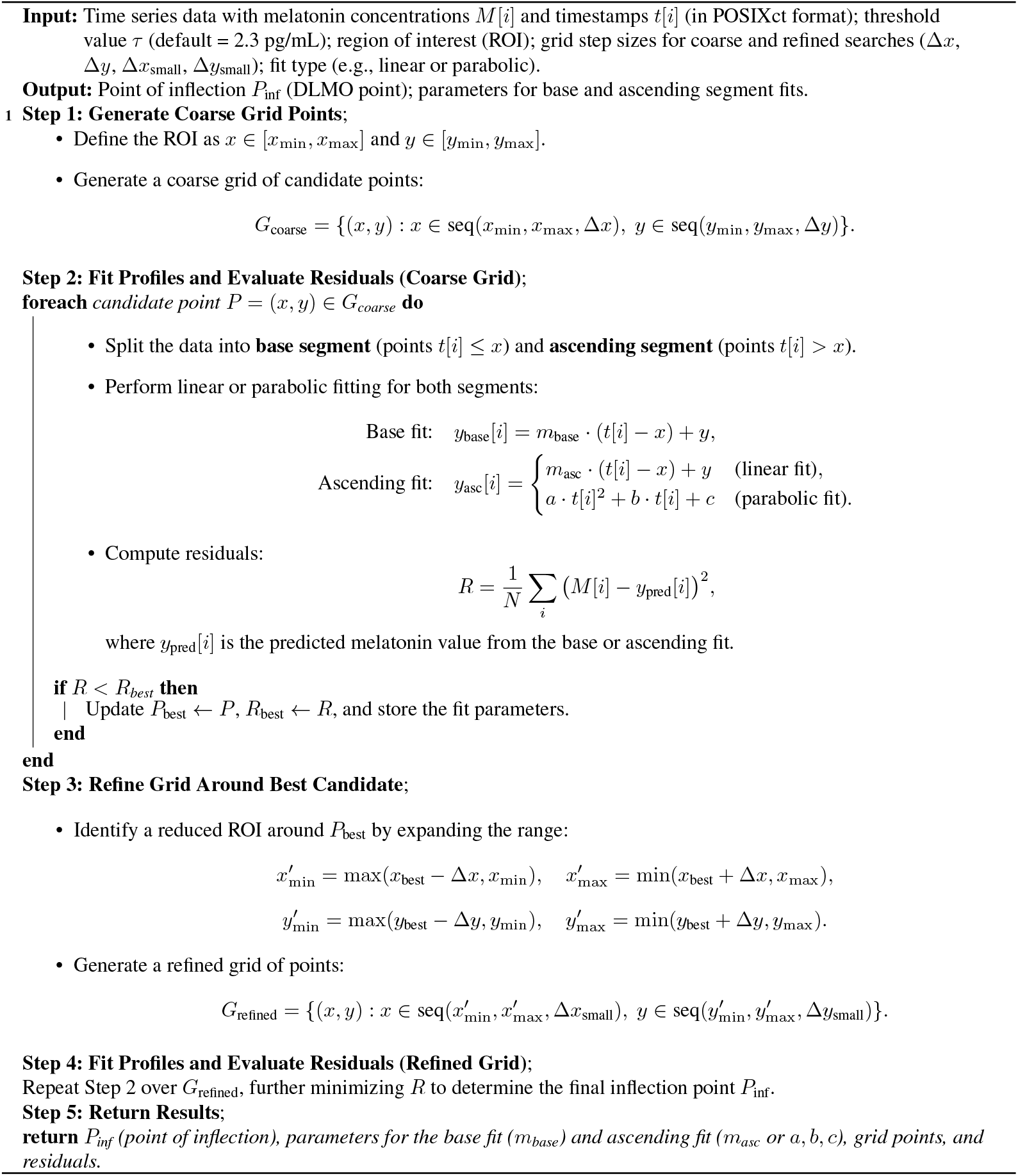

## Appendix B Hockey-stick algorithm parameter rationale

## Appendix C Visualization of outlier cases

## Appendix D Residual heatmap explorer Shiny app

## Appendix E Bootstrapping methods in *dlmoR*

### E.1 Comparison of DLMO estimate bootstrapping methods

### E.2 Example outputs of DLMO estimate variability across bootstrap schemes

## Appendix F Comparison of *dlmoR* and *hockeystickexe* features

## Appendix G Computational runtime of hockey-stick alogrithm analyses

**Table 4.** Wall-clock time and core-hour usage for analyses of the Heinrichs and Spitschan (2025) dataset.

| *Analysis* | *Machine* | *Wall-clock time* | *Core-hours* |
| --- | --- | --- | --- |
| Sampling interval | MacBookPro | 3 hours 11 minutes | 41.38 |
| Threshold | MacBookPro | 1 hours 31 minutes | 19.72 |
| Single-deletion | MacBookPro | 1 hours 31 minutes | 19.72 |
| Multiple deletion | HPC Cluster | 11 hours 46 minutes | 282 |
| Noise | HPC Cluster | 15 hours 15 minutes | 366 |

**Table 5.** Wall-clock time and core-hour usage for analyses of the Blume et al. (2024) dataset.

| *Analysis* | *Machine* | *Wall-clock time* | *Core-hours* |
| --- | --- | --- | --- |
| Sampling interval | MacBookPro | 5 hours 02 minutes | 65.43 |
| Threshold | MacBookPro | 2 hours 44 minutes | 35.53 |
| Single-deletion | MacBookPro | 9 hours 55 minutes | 9.92 |
| Multiple deletion | HPC Cluster | 21 hours 26 minutes | 1372 |
| Noise | HPC Cluster | 27 hours 14 minutes | 1742 |

## Appendix H Analysis summary tables

### H.1 DLMO estimate sensitivity to sampling interval

**Table 6.** Δ DLMO *vs* sampling interval (Heinrichs et al.)

| *interval [min]* | *mean $\Delta$ [h]* | *SD $\Delta$ [h]* | *SE $\Delta$ [h]* | *n* |
| --- | --- | --- | --- | --- |
| 2 | -0.2522 | 0.3540 | 0.0723 | 24 |
| 5 | -0.2470 | 0.3317 | 0.0677 | 24 |
| 10 | -0.1862 | 0.3402 | 0.0694 | 24 |
| 15 | -0.0763 | 0.2077 | 0.0424 | 24 |
| 20 | -0.0899 | 0.2730 | 0.0557 | 24 |
| 30 | 0.0014 | 0.2295 | 0.0468 | 24 |
| 45 | 0.0877 | 0.2759 | 0.0563 | 24 |
| 60 | 0.0931 | 0.3225 | 0.0658 | 24 |
| 75 | -0.0435 | 0.3765 | 0.0769 | 24 |
| 90 | -0.2103 | 0.7073 | 0.1444 | 24 |

**Table 7.** Δ DLMO *vs* sampling interval (Blume et al.)

| *interval [min]* | *mean $\Delta$ [h]* | *SD $\Delta$ [h]* | *SE $\Delta$ [h]* | *n* |
| --- | --- | --- | --- | --- |
| 2 | -0.0501 | 0.2150 | 0.0225 | 91 |
| 5 | -0.0200 | 0.2136 | 0.0225 | 90 |
| 10 | 0.0025 | 0.1820 | 0.0190 | 92 |
| 15 | -0.0250 | 0.1577 | 0.0166 | 90 |
| 20 | -0.0226 | 0.2853 | 0.0297 | 92 |
| 30 | 0.0000 | 0.0000 | 0.0000 | 93 |
| 45 | -0.0520 | 0.4496 | 0.0479 | 88 |
| 60 | -0.1150 | 0.5322 | 0.0564 | 89 |
| 75 | -0.3486 | 0.6837 | 0.0733 | 87 |
| 90 | -0.1637 | 0.7119 | 0.0763 | 87 |

### H.2 DLMO estimate sensitivity to threshold value

**Table 8.**
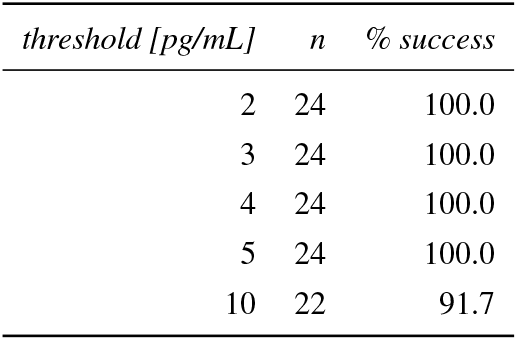
DLMO estimate success per threshold (Heinrichs et al., N=24)

**Table 9.** DLMO estimate success per threshold (Blume et al., N=96)

| *threshold [pg/mL]* | *n* | *% success* |
| --- | --- | --- |
| 2 | 73 | 76.0 |
| 3 | 91 | 94.8 |
| 4 | 93 | 96.9 |
| 5 | 93 | 96.9 |
| 10 | 91 | 94.8 |

**Table 10.**
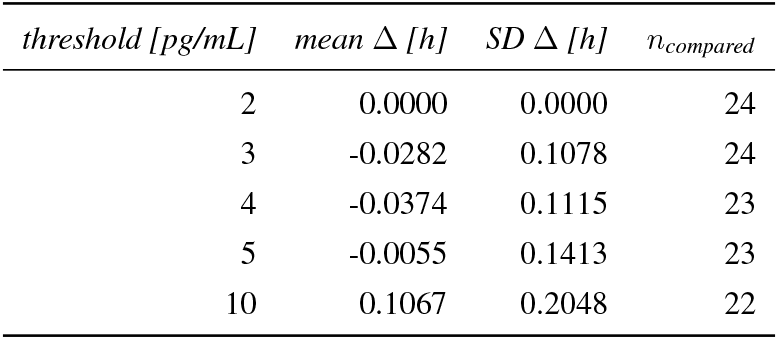
Δ DLMO (relative to threshold = 2) *vs* threshold (Heinrichs et al.)

**Table 11.** Δ DLMO (relative to threshold = 2) *vs* threshold (Blume et al.)

| *threshold [pg/mL]* | *mean $\Delta$ [h]* | *SD $\Delta$ [h]* | *n<sub>compared</sub>* |
| --- | --- | --- | --- |
| 2 | 0.0000 | 0.0000 | 73 |
| 3 | 0.2847 | 0.5856 | 73 |
| 4 | 0.4684 | 0.6782 | 73 |
| 5 | 0.6138 | 0.7473 | 72 |
| 10 | 0.9710 | 1.1041 | 67 |

### H.3 DLMO estimate sensitivity to multi-point deletions

**Table 12.** Δ DLMO *vs* % deletion (Heinrichs et al.)

| % del | mean $\Delta$ [h] | SD $\Delta$ [h] | n | N | % success |
| --- | --- | --- | --- | --- | --- |
| 10 | -0.0409 | 0.4094 | 480 | 480 | 100.0 |
| 20 | -0.0789 | 0.4853 | 480 | 480 | 100.0 |
| 30 | -0.0913 | 0.6187 | 479 | 480 | 99.8 |
| 40 | -0.2384 | 0.8235 | 476 | 480 | 99.2 |
| 50 | -0.3022 | 1.1233 | 466 | 480 | 97.1 |

**Table 13.** Δ DLMO *vs* % deletion (Blume et al.)

| % del | mean $\Delta$ [h] | SD $\Delta$ [h] | n | N | % success |
| --- | --- | --- | --- | --- | --- |
| 10 | -0.0264 | 0.2612 | 1835 | 1920 | 95.6 |
| 20 | -0.0645 | 0.3786 | 1829 | 1920 | 95.3 |
| 30 | -0.1082 | 0.5183 | 1810 | 1920 | 94.3 |
| 40 | -0.1637 | 0.6372 | 1797 | 1920 | 93.6 |
| 50 | -0.2045 | 0.7533 | 1743 | 1920 | 90.8 |

### H.4 DLMO estimate sensitivity to various sources of noise

**Table 14.** Δ DLMO *vs* noise condition, (Heinrichs et al.)

| noise type | mean $\Delta$ [h] | SD $\Delta$ [h] | n | N |
| --- | --- | --- | --- | --- |
| Time only: 5 min | -0.0145 | 0.2137 | 480 | 480 |
| Time only: 10 min | -0.0307 | 0.3641 | 480 | 480 |
| Time only: 20 min | -0.0500 | 0.4940 | 480 | 480 |
| Mel only: 5 min | -0.0303 | 0.3410 | 480 | 480 |
| Mel + time: 10 min | -0.0396 | 0.4363 | 480 | 480 |

**Table 15.**
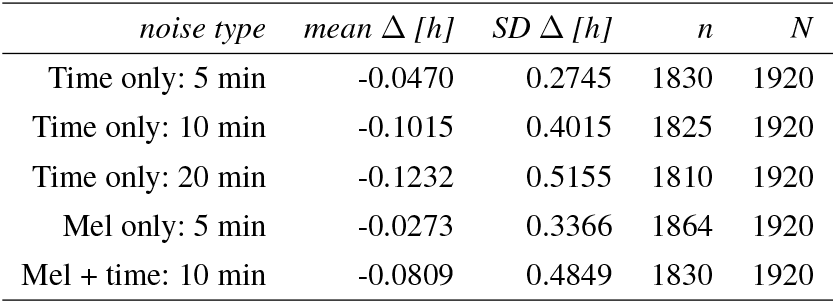
Δ DLMO *vs* noise condition (Blume et al.)

| noise type | mean $\Delta$ [h] | SD $\Delta$ [h] | n | N |
| --- | --- | --- | --- | --- |
| Time only: 5 min | -0.0470 | 0.2745 | 1830 | 1920 |
| Time only: 10 min | -0.1015 | 0.4015 | 1825 | 1920 |
| Time only: 20 min | -0.1232 | 0.5155 | 1810 | 1920 |
| Mel only: 5 min | -0.0273 | 0.3366 | 1864 | 1920 |
| Mel + time: 10 min | -0.0809 | 0.4849 | 1830 | 1920 |

## Appendix I Example profiles for multiple deletion analysis

## Appendix J Guidance on choosing a threshold value for DLMO estimation

Selecting an appropriate threshold is a critical step in estimating dim light melatonin onset (DLMO) using the hockey-stick algorithm. While there is no universally applicable threshold, we recommend the following pragmatic and transparent approach:

1. **Start with a standard threshold** Begin with a fixed, commonly used threshold (e.g., 3 pg/mL) that has been demonstrated to yield reliable DLMO estimates across a wide range of individuals. This approach maximizes consistency and facilitates comparison across studies.
2. **Identify problematic profiles** For some participants, the standard threshold may fail to yield a valid DLMO estimate. This can occur when the entire melatonin profile remains either entirely below (sub-threshold) or above (supra-threshold) the chosen value, or when the melatonin rise is too shallow or too steep to clearly define an onset point. These profiles should be flagged for individual review and may require an adaptive or individualized thresholding approach.
3. **Use a data-driven alternative for flagged cases** In such instances, a statistical threshold (e.g., the mean plus 2 standard deviations of the first 3 samples) (see Voultsios et al. (1997); Molina and Burgess (2011); Kennaway (2023); Murray et al. (2024), may provide a more appropriate and biologically meaningful estimate. Any deviation from the standard method should be pre-specified or decided by consensus between two independent raters to ensure objectivity.
4. **Report transparently** Clearly document the thresholding strategy in the methods section. For transparency and reproducibility, we recommend including a supplementary table listing the threshold value used for each participant, especially in cases where an individualized threshold was applied.

This structured approach enables consistency across the majority of participants while allowing flexibility for challenging cases. It also promotes transparency and interpretability in reporting DLMO estimation procedures.

## Footnotes

* The description of this portion of the algorithm in the publication by Danilenko et al. (2014) differed from the natural-language explanation provided in personal correspondence, in regards to the use of weighting for the L_2_-norms of the fits based on base labels. In our dltnoR implementation, we chose not to apply any weighting, and after conducting multiple trials, we found no evidence that weighting improved the results.

## References

(2008) R news: The newsletter of the r project. URL https://www.r-project.org/doc/Rnews/Rnews_2008-1.pdf. The Newsletter of the R Project.

Benloucif S, Burgess HJ, Klerman EB, Lewy AJ, Middleton B, Murphy PJ, Parry BL and Revell VL (2008) Measuring melatonin in humans. Journal of Clinical Sleep Medicine 4(1): 66–69.

Blume C, Cajochen C, Schöllhorn I, Slawik HC and Spitschan M (2024) Effects of calibrated blue–yellow changes in light on the human circadian clock. Nature Human Behaviour 8(3): 590–605.

Crowley SJ, Poole E, Adams J and Eastman CI (2024) Extending weeknight sleep duration in late-sleeping adolescents using morning bright light on weekends: a 3-week maintenance study. Sleep Advances 5(1): zpae065.

Danilenko K and Verevkin EG (2020) Hockey-stick software. URL https://neuronm.ru/R&D/hockey-stick.

Danilenko KV, Verevkin EG, Antyufeev VS, Wirz-Justice A and Cajochen C (2014) The hockey-stick method to estimate evening dim light melatonin onset (DLMO) in humans. Chronobiology International 31(3): 349–355.

Fasola S, Muggeo VM and Küchenhoff H (2018) A heuristic, iterative algorithm for change-point detection in abrupt change models. Computational Statistics 33(2): 997–1015.

Glacet R, Reynaud E, Robin-Choteau L, Reix N, Hugueny L, Ruppert E, Geoffroy PA, Kilic-Huck Ü, Comtet H and Bourgin P (2023) A comparison of four methods to estimate dim light melatonin onset: a repeatability and agreement study. Chronobiology International 40(2): 123–131.

Harrison S, Dasgupta A, Waldman S, Henderson A and Lovell C (2021) How reproducible should research software be? URL https://zenodo.org/records/4761867.

Heinrichs HS and Spitschan M (2025) Within-subjects ultra-short sleep-wake protocol for characterising circadian variations in retinal function. PloS one 20(5): e0300405.

Keijzer H, Smits MG, Duffy JF and Curfs LM (2014) Why the dim light melatonin onset (DLMO) should be measured before treatment of patients with circadian rhythm sleep disorders. Sleep Medicine Reviews 18(4): 333–339.

Kennaway DJ (2023) The dim light melatonin onset across ages, methodologies, and sex and its relationship with morningness/eveningness. Sleep 46(5): zsad033.

Lee G, Bacon S, Bush I, Fortunato L, Gavaghan D, Lestang T, Morton C, Robinson M, Rocca-Serra P, Sansone SA et al. (2021) Barely sufficient practices in scientific computing. Patterns 2(2).

Molina TA and Burgess HJ (2011) Calculating the dim light melatonin onset: the impact of threshold and sampling rate. Chronobiology international 28(8): 714–718.

Muggeo VM (2003) Estimating regression models with unknown break-points. Statistics in medicine 22(19): 3055–3071.

Muggeo VM (2017) Interval estimation for the breakpoint in segmented regression: a smoothed score-based approach. Australian & New Zealand Journal of Statistics 59(3): 311–322.

Murray JM, Stone JE, Abbott SM, Bjorvatn B, Burgess HJ, Cajochen C, Dekker JJ, Duffy JF, Epstein LJ, Garbazza C et al. (2024) A protocol to determine circadian phase by athome salivary dim light melatonin onset assessment. Journal of Pineal Research 76(5): e12994.

Pan Y (2022) roptim: General Purpose Optimization in R using C++. DOI:10.32614/CRAN.package.roptim. R package version 0.1.6.

Pandi-Perumal SR, Smits M, Spence W, Srinivasan V, Cardinali DP, Lowe AD and Kayumov L (2007) Dim light melatonin onset (DLMO): a tool for the analysis of circadian phase in human sleep and chronobiological disorders. Progress in Neuro-Psychopharmacology and Biological Psychiatry 31(1): 1–11.

Pedersen EJ, Miller DL, Simpson GL and Ross NE (2019) Hierarchical generalized additive models in ecology: An introduction with mgcv. PeerJ 7: e6876. DOI:10.7717/peerj.6876.

Putilov AA (2023) Obituary for Dr. Konstantin Danilenko (19.03.1962–18.01.2023). Clocks & Sleep 5(1): 94–97. DOI: 10.3390/clockssleep5010010.

R Core Team (2024) R: A Language and Environment for Statistical Computing. R Foundation for Statistical Computing, Vienna, Austria. URL https://www.R-project.org/.

Rasmussen CE and Williams CK (2006) Gaussian Processes for Machine Learning. Cambridge, MA: MIT Press. URL http://www.gaussianprocess.org/gpml/.

Voultsios A, Kennaway DJ and Dawson D (1997) Salivary melatonin as a circadian phase marker: validation and comparison to plasma melatonin. Journal of biological rhythms 12(5): 457–466.

Wickham H (2016) ggplot2: Elegant Graphics for Data Analysis. Springer-Verlag New York. ISBN 978-3-319-24277-4. URL https://ggplot2.tidyverse.org.

Wickham H, Averick M, Bryan J, Chang W, McGowan L, François R, Grolemund G, Hayes A, Henry L, Hester J, Kuhn M, Pedersen TL, Miller E, Bache SM, Müller K, Ooms J, Robinson D, Seidel D, Spinu V, Takahashi K, Dunnington D, Wilke C, Woo K and Yutani H (2019) Welcome to the tidyverse. Journal of Open Source Software 4(43): 1686. DOI:10.21105/joss.01686. URL https://doi.org/10.21105/joss.01686.

Wickham H, François R, Henry L and Müller K (2023) dplyr: A Grammar of Data Manipulation. URL https://dplyr.tidyverse.org. R package version 1.1.2.

Wilson G, Bryan J, Cranston K, Kitzes J, Nederbragt L and Teal TK (2017) Good enough practices in scientific computing. PLoS Computational Biology 13(6): e1005510.

Wright KP, McHill AW, Birks BR, Griffin BR, Rusterholz T and Chinoy ED (2013) Entrainment of the human circadian clock to the natural light-dark cycle. Current biology 23(16): 1554–1558.

Zerbini G, Winnebeck EC and Merrow M (2021) Weekly, seasonal, and chronotype-dependent variation of dim-light melatonin onset. Journal of Pineal Research 70(3): e12723.

