## Supplementary figures and images for "*dlmoR:* An open-source R package for the dim-light melatonin onset (DLMO) hockey-stick method"

### Graphical abstract

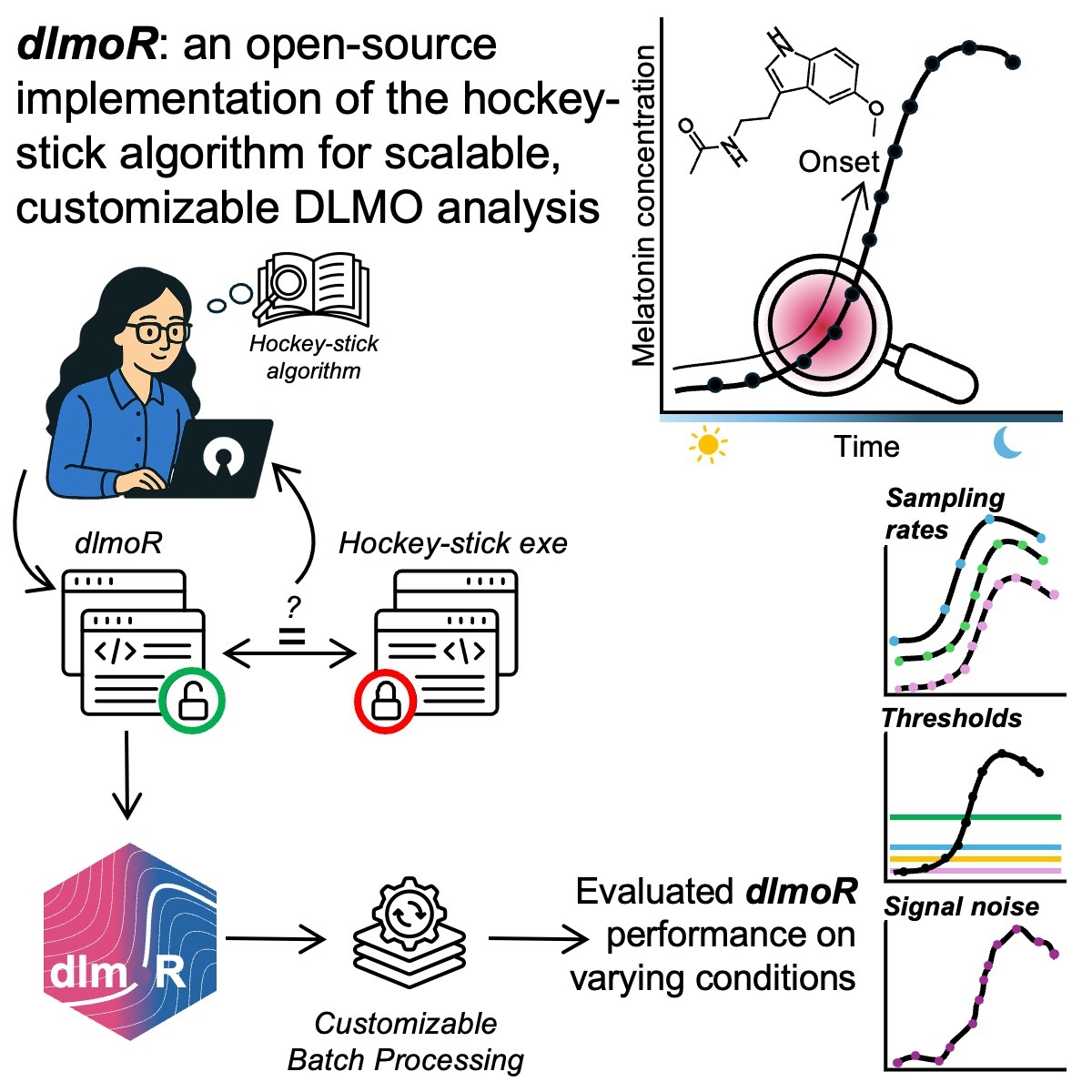
